# Decoding Specificity of Cyanobacterial MysDs in Mycosporine-Like Amino Acid Biosynthesis through Heterologous Expression in *Saccharomyces cerevisiae*

**DOI:** 10.1101/2024.09.14.613006

**Authors:** Xiaoyou Zheng, Peifeng Xie, Andrew Chen Cai, Yuze Jiang, Sirui Huang, Xiaochong Ma, Honghao Su, Boxiang Wang

**Affiliations:** Churchill College, University of Cambridge, Storey’s Way, Cambridge, United Kingdom, CB3 0DS; LINK SPIDER Co., Ltd., 11 Langshan Rd, Nanshan District, Shenzhen, China, 518000; Thurgood Marshall College, University of California, San Diego, 9500 Gilman Dr., La Jolla, California, United States; Earlham Institute, Norwich Research Park, Norwich, United Kingdom, NR4 7UZ

## Abstract

Mycosporine-like amino acids (MAAs) are potent natural UV-protectants, but their industrial production is hindered by efficiency and sustainability issues of large-scale extraction of their native hosts. Heterologous expression of MAA biosynthesis pathway genes in chassis organisms provides a promising alternative route, though the substrate promiscuity of the ATP-grasp ligase MysD complicates the biosynthesis of specific MAAs. In this study, we developed a *Saccharomyces cerevisiae* strain with enhanced capacity of producing mycosporine-glycine (MG), through genomic expression of biosynthesis pathway genes and knockout of competing pathway genes. This strain serves as an efficient MysD expression platform, which converts MG into shinorine and porphyra-334. Through structural modelling, site-directed mutagenesis and mutant characterization, we identified two residues on the omega-loop of MysD involved in determining product specificity. We further characterized the product specificity of 20 MysDs from diverse cyanobacterial lineages and confirmed the residue pattern-product specificity correlation. Our findings provide guidance for screening, selecting, and designing novel MysDs for industrial-scale MAA production through heterologous expression.

## Introduction

Prolonged exposure to ultraviolet radiation (UVR) from the sun is associated with increased risk of erythema, oxidative injury, photoaging and skin cancer, creating a need of effective sunscreens. ^1^ Mycosporine-like amino acids (MAAs), photo-protectants capable of absorbing both UVA (315-400nm) and UVB (280-315nm), are found in diverse UV-adapted marine and terrestrial organisms, including several lineages of actinobacteria, cyanobacteria and diatoms.^2–5^ Compared to conventional sunscreens with zinc oxide and oxybenzone content, MAAs demonstrate higher biocompatibility and less environmental impact, which make them promising candidates of natural sunscreens and attractive for industrial production.^6,7^ Traditional method of MAA production involves large-scale harvesting, processing and extraction of marine algae, which suffers from low overall yield and product purity and raises the issue of profitability and sustainability.^8^ As an alternative means, heterologous production in chassis microorganisms including *Escherichia coli* and *Saccharomyces cerevisiae* by engineering metabolic pathways demonstrates the potential of more efficient and specific production of industrially valuable natural product molecules, including MAAs.^9,10^ Hence, to engineer MAA production in heterologous hosts, knowledge of MAA biosynthesis pathway is needed.

Biosynthetic gene clusters (BGCs) of MAA have been discovered in several cyanobacteria species and most of them contain three conserved genes, including *mysA* which encodes a DDG synthase (DDGS), *mysB* which encodes a SAM-dependent O-methyltransferase (O-MT) and *mysC* which encodes an ATP-grasp enzyme.^11^ MysA and MysB catalyze the stepwise conversion of sedoheptulose 7-phosphate (S7P) from pentose phosphate pathway (PPP) into 4-deoxygadusol (4-DG), the scaffold of all MAAs, while MysC subsequently catalyzes the addition of glycine to 4-DG and gives rise to mycosporine-glycine (MG) (Figure 1A). In several cyanobacteria strains, BGCs of MAA also include *mysD*, which encodes a homolog of D-alanine-D-alanine ligase.^11,12^ MysD is capable of catalyzing the conjugation of an amino acid to MG, which gives rise to different types of di-substituted MAAs, including shinorine (MG + L-Serine) and porphyra-334 (MG + L-threonine) (Figure 1A).^13,14^

**Figure 1.**
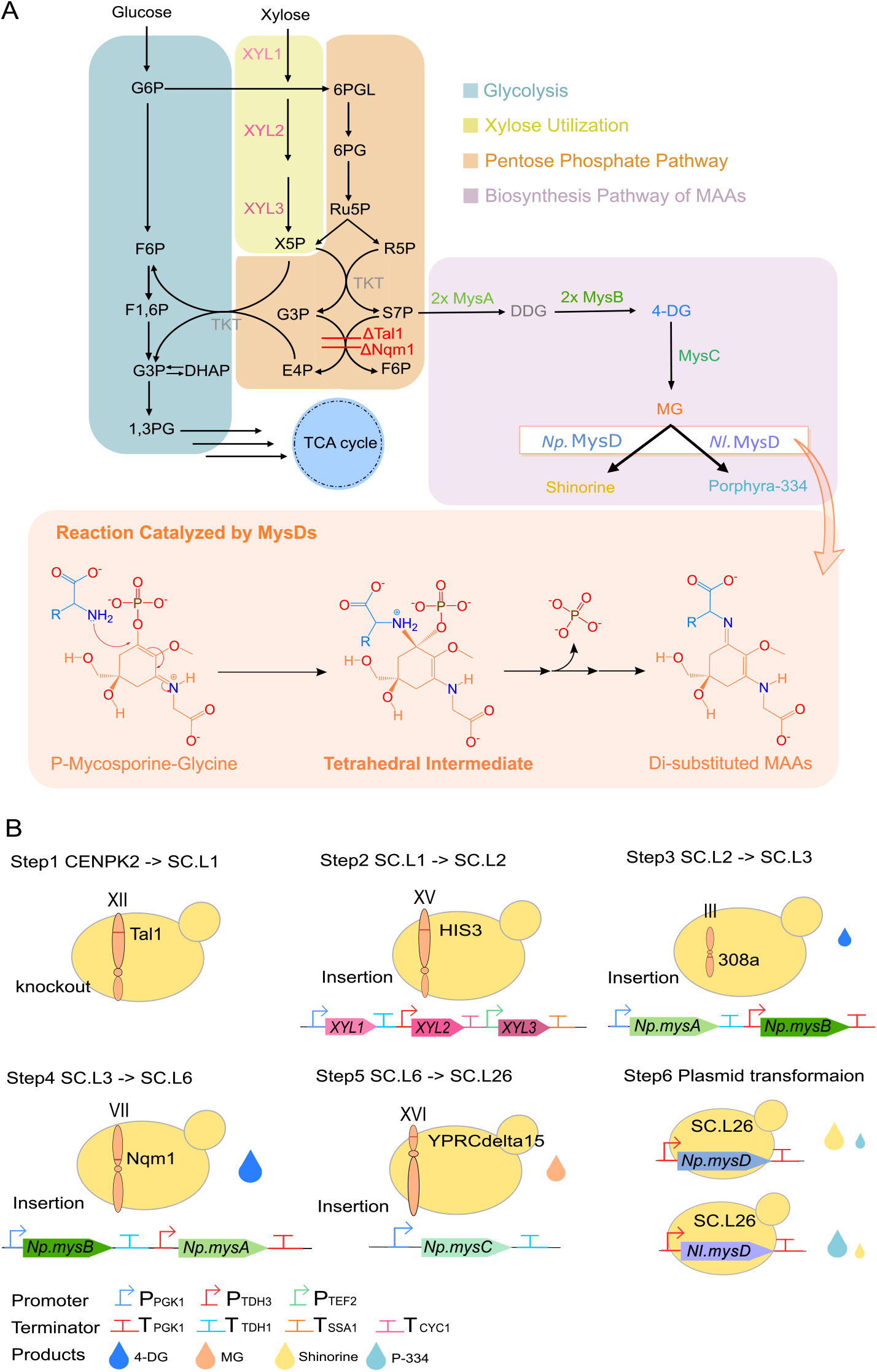
(A) Engineered metabolic pathway for the biosynthesis of disubstituted MAAs. The knockout of the transaldolase gene *Tal1* and its paralog *Nqm1* (Red) limits the conversion of S7P into the unwanted F6P. The introduction of the xylose-utilizing pathway *xyl1, 2*, and *3* (Magenta) enhances S7P production by directing xylose into PPP. The double knock-in of *mysA* and *mysB* and the introduction of *mysC* (Green) achieve the biosynthesis of MG. *mysDs* including *Np*.*mysD and Nl*.*mysD* (Blue and Purple) directs the synthesis of disubstituted MAAs including shinorine and porphyra-334 (Yellow and Blue). The reaction mechanism from MG to di-substituted MAAs is shown. The tetrahedral intermediate (middle) was used for molecular docking. Abbreviations of metabolites are provided in the glossary of supplementary materials. (B) Step-wise yeast engineering for MAA production. The knock-in of the second copy of *mysA* and *mysB* and the deletion of the *Nqm1* gene were simultaneously accomplished in step4.

According to previous analysis on the mode-of-action of D-alanine-D-alanine ligases and predicted catalytic mechanism of MysDs,^12,15,16^ the reaction from MG to di-substituted MAAs begins with the phosphorylation of the C1 position of MG. This is followed by a nucleophilic attack from the amine moiety of the amino acid substrate on the same position (C1). These actions result in the formation of a tetrahedral, phosphate-containing intermediate (Figure 1A, Figure S5). The reaction is driven to completion by the dissociation of the phosphate group, followed by the release of MAA product and ADP from MysD.

All characterized MysDs demonstrate substrate and product promiscuity and produce more than one type of di-substituted MAAs, mostly a mixture of shinorine and porphyra-334.^14^ However, different MysDs produce shinorine and porphyra-334 in different ratios.^12–14,17,18^ Nonetheless, except for Kim et al., 2023 which correlated product specificity with omega loop, few studies have explored the molecular basis of the difference in product specificity amongst MysDs.^14^ Hence, a thorough understanding of the molecular basis of MysD product specificity needs to be developed to inform the engineering of MysD for more efficient and specific di-substituted MAA production.

In this study, we explored means of enhancing efficiency and specificity of the production of di-substituted MAAs shinorine and porphyra-334. Through heterologous pathway expression and knockdown of competing pathways, we developed a yeast strain capable of producing MG. We used structural modelling and site-directed mutagenesis and identified two critical amino acid residues associated with product specificity of MysD. We validated the importance of the two residues in product specificity determination through sequence-informed selection and characterization of previously-uncharacterized MysDs.

## Results and Discussion

### Engineering Yeast Strains for MG production

MG is the common precursor of shinorine and porphyra-334, two di-substituted MAAs of interest (Figure 1A). Therefore, a *S. cerevisiae* strain efficient at producing MG needs to be engineered to pave way for shinorine and porphyra-334 production. Based on the strategy of Park et al., 2019,^18^ we first sequentially knocked out *Tal1* and engineered xylose-utilization pathway by inserting three *Scheffersomyces stipitis* xylose-utilizing genes (*Xyl1, Xyl2* and *Xyl3*) in *HIS3* locus to enhance the production of S7P, a metabolite in pentose phosphate pathway that serves as the ultimate precursor of MAA biosynthesis pathway (Figure 1A, 1B), in *S. cerevisiae* strain CEN-PK, giving rise to strain SC.L2.

To enable the conversion of S7P to 4-DG, we sequentially made strain SC.L3 and SC.L6 by inserting either just one copy of *Np*.*mysA* and *Np*.*mysB* from *Nostoc punctiforme* into 308 locus or another copy of them into *Nqm1* locus, which encodes a transaldolase that diverts S7P flux from MAA biosynthesis pathway (Figure 1A, 1B). To confirm their 4-DG production capability, both strains were cultured and extracted and LC-HRMS analysis was performed on their extracts (Figure 2A). In the chromatogram, peak B, whose MS fragmentation pattern is consistent with 4-DG, was observed in the extracts of SC.L3 and SC.L6 but not SC.L2. Therefore, we validated that both strains are capable of producing 4-DG. Through peak area comparison, we observed a significant increase in 4-DG production in SC.L6 compared to SC.L3 (p<0.05, one-way ANOVA) with a 3.00-fold difference (Figure 2B). SC.L6 was therefore used for engineering of further steps.

**Figure 2.**
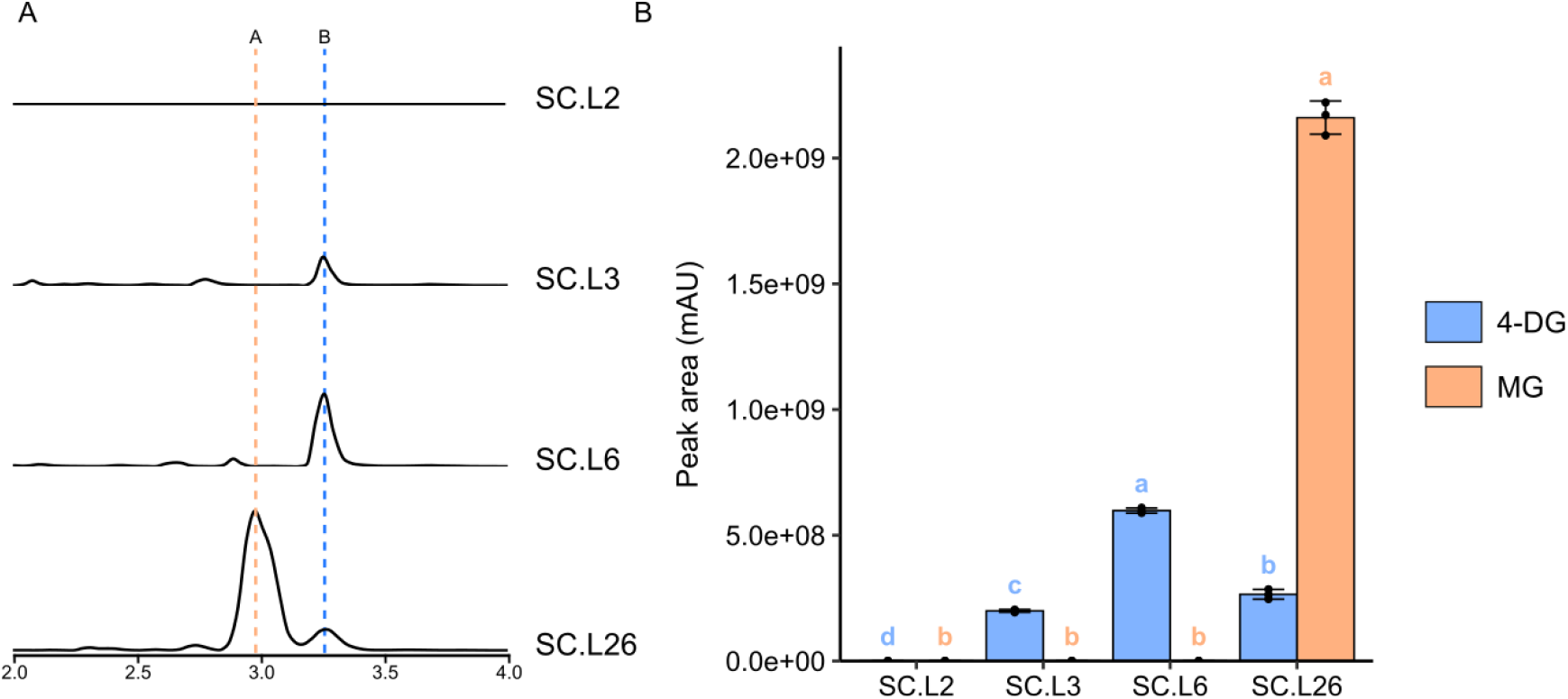
Engineering *S. cerevisiae* for MG production. (A) Extracted ion chromatograms (EICs) of LC-HRMS analysis of ions with m/z= 188.08-189.08 and 245.10-246.10 in SC.L2, SC.L3, SC.L6 and SC.L26 samples. Peak B’s retention time is 3.24 min and peak A’s retention time is 2.93 min and their identities were confirmed as 4-DG and MG through MS analysis. The height each chromatogram is 2.5×10E^8^ arbitrary unit. (B) 4-DG and MG peak area comparison amongst these yeast strains. Heights of the bars represent their means and the error bars represent their standard deviations (n=3 biological replicates). One-way ANOVA was performed to compare the amount of 4-DG and MG separately amongst different samples and two samples that do not share the same letter are significantly different from each other (p<0.05).

To complete the engineering of MG production pathway, we inserted *Np*.*mysC* into locus *YPRCdelta15* of SC.L6 to form strain SC.L26 (Figure 1A, 1B). SC.L26 was also cultured and extracted and LC-HRMS was performed on the extract (Figure 2A). In the chromatogram, both peak B and peak A, a new peak not found in any other analyzed strains, were observed in SC.L26’s extract. Peak A was later confirmed as MG through MS pattern comparison.

### Identification of two specificity-determining residues on the omega-loop of MysD

Stepwise pathway engineering enabled the development of a *S. cerevisiae* strain (SC.L26) capable of producing MG. To convert MG into shinorine and porphyra-334, further engineering of SC.L26 by expressing MysD is needed. All characterized MysD homologs show substrate and product promiscuity and produce both shinorine and porphyra-334 simultaneously.^12–14,17,18^ Nevertheless, substrate preference and product specificity vary amongst them. Hence, to engineer product ratio for more specific production of one type of MAA, an understanding of structure-product specificity relationship needs to be developed.

Difference in product specificity is generally associated with difference in active site residues. To look for that, we first selected MysDs from *Nostoc punctiforme* PCC 73102 (*Np*.MysD) and from *Nostoc linckia* NIES-25 (*Nl*.MysD) as exemplars of shinorine-preferred producer and porphyra-334-preferred producer respectively (Figure 1A). We acquired their structural models through ColabFold and found high structural homology between each other (RMSD=0.795) and with *E*.*coli* D-Alanine-D-Alanine ligase’s crystal structure^16^ (PDB ID: 2DLN) (RMSD=1.901 for *Np*.MysD and 1.794 for *Nl*.MysD) (Figure 3A, Figure S1). We predicted the active site pockets of MysDs based on 2DLN and docked the tetrahedral intermediate (MG-P-Ser) into their pockets (Figure 3A). We hypothesize that residues less than 8 angstroms from the docked intermediates that differ between *Np*.MysD and *Nl*.MysD are involved in substrate recognition and hence product specificity determination.

**Figure 3.**
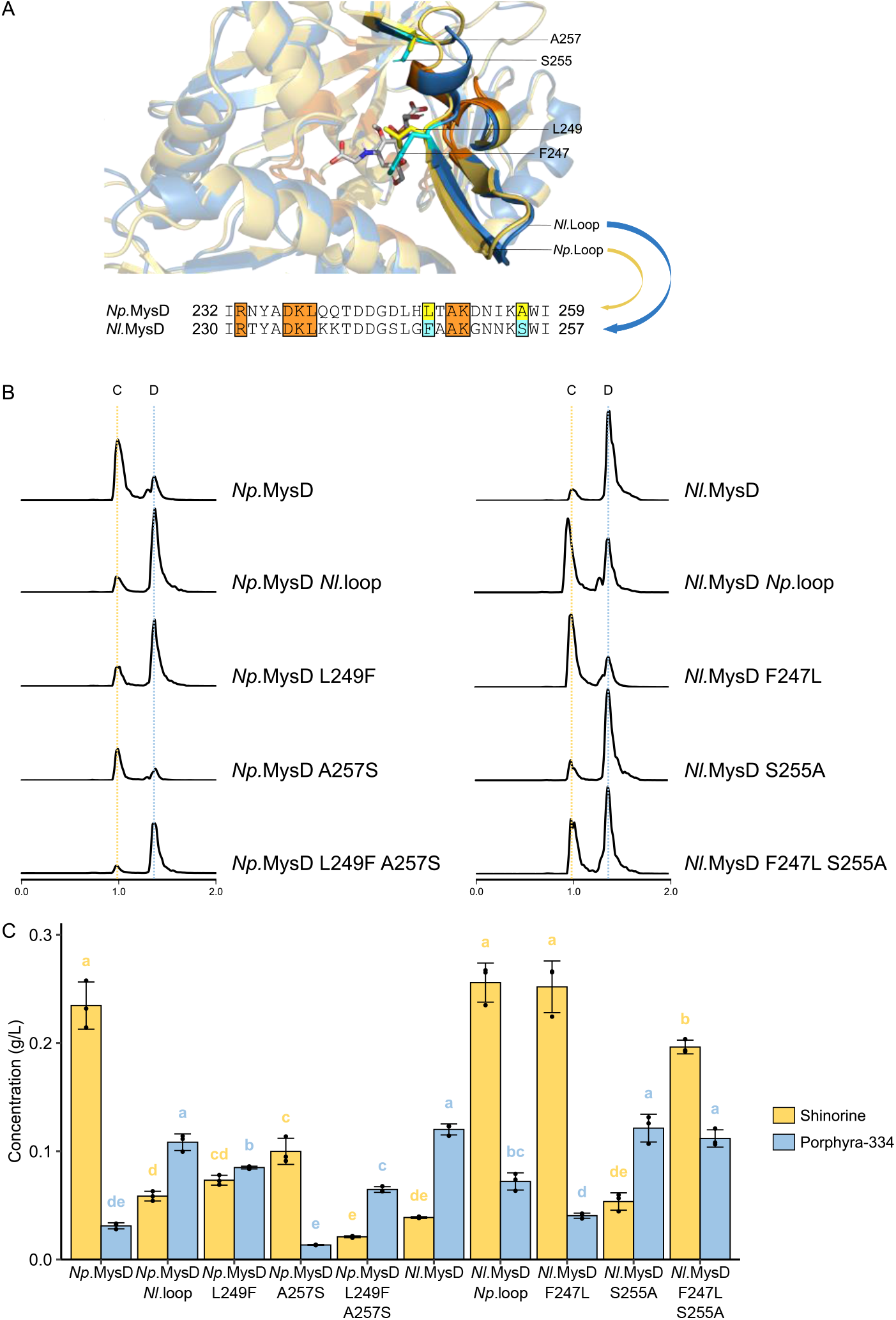
(A) Predicted structural models of *Np*.MysD (blue) and *Nl*.MysD (yellow) with a reaction intermediate (M-Gly-P-Ser) docked to active sites and an alignment of their omega-loop sequences. The identical sequences within 5 angstroms from the intermediate are colored orange. The two critical aa residues are highlighted with yellow (*Np*.MysD) or cyan (*Nl*.MysD). (B) EICs of LC-HRMS analysis of ions with m/z= 332.50-333.50 and 346.50-347.50 in SC.L26 expressing native and mutant *Np*.MysD and *Nl*.MysD. Peak C’s retention time is 1.00 min and peak D’s retention time is 1.38 min and their identities were confirmed as shinorine and porphyra-334 through MS analysis. The height of each chromatogram is 1.2×10E^9^ arbitrary unit. (C) Amount of shinorine and porphyra-334 produced by native and mutant *Np*.MysD and *Nl*.MysD. Concentrations were calculated by adjusting raw peak areas with their respective response factors. Heights of the bars represent their means and the error bars represent their standard deviations (n=3 biological replicates). One-way ANOVA was performed to compare the amount of shinorine and porphyra-334 separately amongst different samples and two samples that do not share the same letter are significantly different from each other (p<0.05).

All active site residues that differ between *Np*.MysD and *Nl*.MysD are clustered in a 26-amino acid long segment located on the 45-amino acid product-specificity determining omega-loop discovered by Kim et al., 2023 (Figure 3A, Figure S2).^14^ We performed loop substitution on both *Np*.MysD and *Nl*.MysD and expressed wild-type and loop-substituted *Np*.MysDs and *Nl*.MysDs in SC.L26. By analyzing their metabolite contents, we found that *Np*.MysD-*Nl*.Loop produces predominantly porphyra-334 (peak D) while *Nl*.MysD-*Np*.Loop produces predominantly shinorine (peak C), both of which display a reverse in dominant product compared to their corresponding wild-types (Figure 3B), which is corroborates on the previous study.^14^ Peak area analysis demonstrated a significant reduction in dominant product and a significant increase in non-dominant product in the loop-substitution mutants compared to their corresponding wild-types (Figure 3B). However, we observed a reduction in total activity in *Np*.MysD-*Nl*.Loop while an enhancement in total activity in *Nl*.MysD-*Np*.Loop (Figure S3).

To look for a more precise product-specificity determination pattern, we narrowed our scope to residues within 5 angstroms from the intermediate and found only two residues (249 and 257 in *Np*.MysD) that differ between *Np*.MysD and *Nl*.MysD (Figure 3A, 3B). To validate the role of these two residues, we made reciprocal mutants for *Np*.MysD and *Nl*.MysD by swapping either one residue or both of them and expressed them in SC.L26.

For both *Np*.MysD and *Nl*.MysD, a switch in dominant product was observed in L249F / F247L single mutants and L249F-A257S double mutant, with *Np*.MysD mutants producing predominantly porphyra-334 (peak C) and *Nl*.MysD mutants producing predominantly shinorine (peak D) (Figure 3B). However, the switch was not observed in *Np*.MysD-A257S, *Nl*.MysD-S255A and *Nl*.MysD-F247L-S255A.

The response factor of the 0.1 mg/ml porphyra-334 standard sample is 2.7 times greater than that of the 0.1 mg/ml shinorine standard sample. This substantial difference indicates that analyzing raw LC chromatograms alone is insufficient to determine the product specificity. Consequently, we calculated the concentrations of shinorine and porphyra-334 in all samples by extracting peak areas with their response factors adjusted (Figure 3C).

In peak area analysis, we found a significant reduction in shinorine production in all *Np*.MysD mutants and a significant increase in porphyra-334 production in all mutants except *Np*.MysD-A257S (Figure 3C), suggesting that the two residues picked are involved in determining product specificity in *Np*.MysD. *Np*.MysD-L249F-A257S double mutant shows higher porphyra-334-to-shinorine ratio than both single mutants, suggesting potential concerted action between the two residues. However, a significant decline in overall activity was observed in all *Np*.MysD mutants (Figure S3).

For *Nl*.MysD, a significant increase in shinorine production was observed in both *Nl*.MysD-F247L and *Nl*.MysD-F247L-S255A mutants while a significant reduction in porphyra-334 production was observed only in *Nl*.MysD-F247L mutant (Figure 3C). No significant difference in shinorine or porphyra-334 production was observed in *Nl*.MysD-S255A mutant compared to wild-type. *Nl*.MysD-F247L single mutant shows higher shinorine-to-porphyra-334 ratio than other two mutants, suggesting that S255A mutation might reduce the effect of F247L mutation. In addition, we observed a significant boost in activity in both *Nl*.MysD-F247L and *Nl*.MysD-F247L-S255A mutants compared to wild-type (Figure S3).

The mutagenesis results narrowed product specificity determinant of MysD down to the identities of two residues on omega loop. Based on metabolite content produced by wild-type and mutant *Np*.MysD and *Nl*.MysD, we showed a pattern in which MysD preferentially produces shinorine if the two residues are L-A or L-S, while MysD preferentially produces porphyra-334 if the two residues are F-S or F-A. The pattern of these two residues could provide guidance in screening for more efficient and specific shinorine or porphyra-334 producers.

### Characterization of product specificity of various cyanobacterial MysD orthologs selected based on specificity-determining residues

To further validate the link between the identity of these two residues on omega-loop and product specificity, we set out to characterize MysD homologs in other species with varied residue pattern. We interrogated OrthoDB and NCBI database with BLASTp using *Np*.MysD and *Nl*.MysD as query sequences, performed multiple sequence alignment of the hits and chose 20 more sequences to characterize based on the identities of the two product-specificity determining residues (Figure 4A). We acquired their coding sequences through gene synthesis and subsequently expressed them in yeast strain SC.L26. LC-HRMS followed by peak area analysis were applied to evaluate the amount of shinorine and porphyra-334 produced by each MysD. Response factors for shinorine and porphyra-334 were similar in this round of analysis and no response factor adjustment was applied.

**Figure 4.**
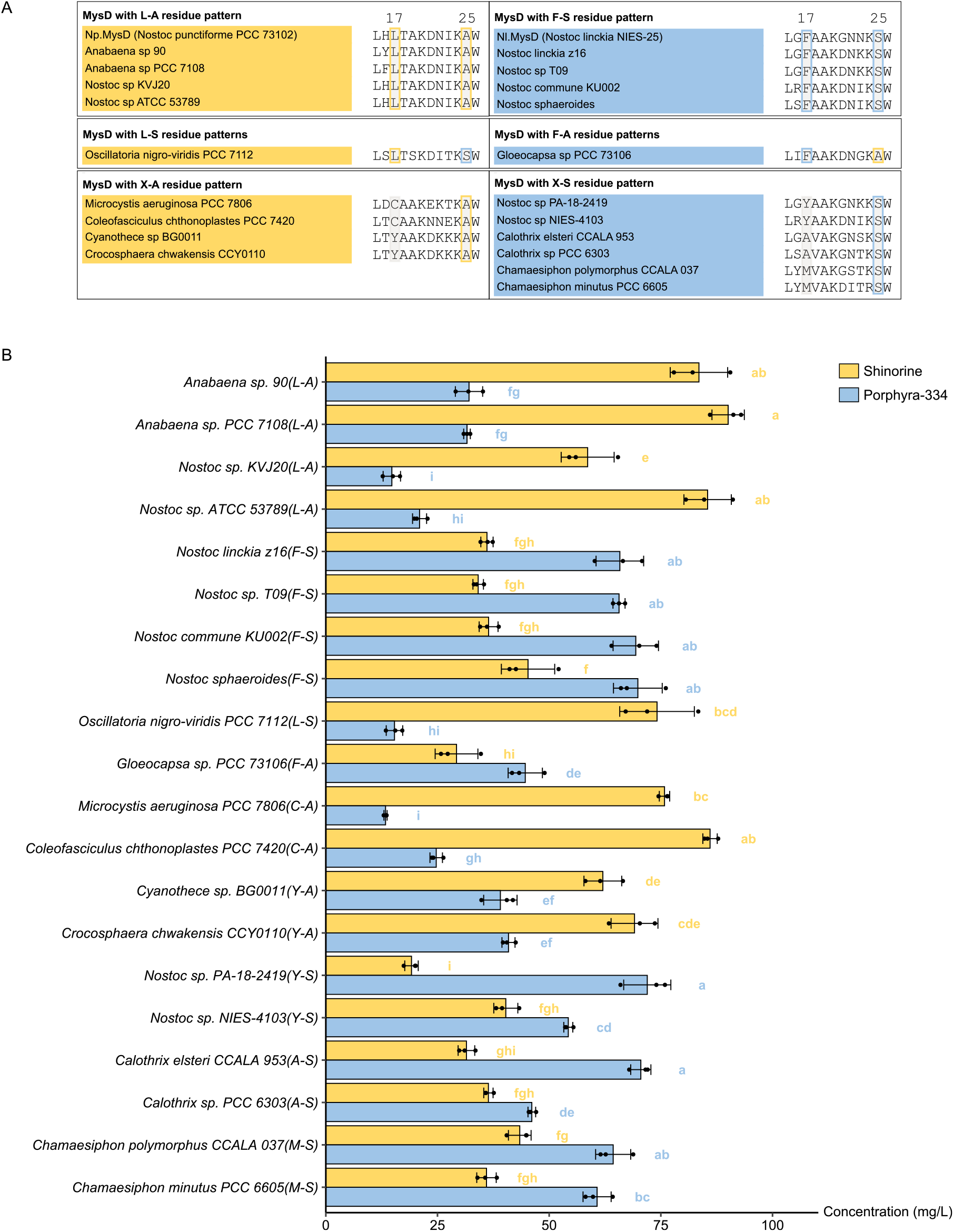
(A) Classification of MysD sequences characterized in this study based on the pattern of two key residues. Sequences hypothesized to produce more shinorine are labelled in yellow while and those hypothesized to produce more porphyra-334 are labelled in blue. Two key residues are highlighted by rectangles with grey background color. (B) Shinorine and porphyra-334 production from LC-HRMS peak area analysis by yeast strains expressing selected MysDs. Lengths of the bars represent their means and the error bars represent their standard deviations (n=3 biological replicates). One-way ANOVA was performed to compare the amount of shinorine and porphyra-334 separately amongst different samples and two samples that do not share the same letter are significantly different from each other (p<0.05).

We first hypothesized that all MysDs with L-A pattern (same as wild-type *Np*.MysD) preferentially produce shinorine while all MysDs with F-S pattern (same as wild-type *Nl*.MysD) preferentially produce porphyra-334. Therefore, we chose four MysDs with L-A pattern (*Anabaena* sp. 90, *Anabaena* sp. PCC 7108, *Nostoc* sp. KVJ20, *Nostoc* sp. ATCC 53789) and four MysDs with F-S patterns (*Nostoc linckia* z16, *Nostoc* sp. T09, *Nostoc commune* KU002, *Nostoc sphaeroides*) for characterization. Consistent with our hypothesis, all MysDs selected with L-A pattern produce more shinorine while all MysDs selected with F-S pattern produce more porphyra-334 when expressed in SC.L26 (Figure 4B, Figure S9-S16).

In the mutagenesis experiment, we found that *Nl*.MysD-F247L mutant which shows L-S pattern preferentially produces shinorine while *Np*.MysD-L249F mutant which shows F-A pattern preferentially produces porphyra-334. To test if the pattern-product specificity relationship also holds true for natural MysDs, we included one MysD with L-S pattern (*Oscillatoria nigro-viridis* PCC 7112) and one MysD with F-A pattern (*Gloeocapsa* sp. PCC 73106) for characterization. When expressed in SC.L26, MysD with L-S pattern produces more shinorine while MysD with F-A pattern produces more porphyra-334, which is consistent with our expectation (Figure 4B, Figure S17-S18).

Finally, we set out to look at the product specificity of MysDs with other patterns at the two residues. Given MysD with L-A pattern and previously characterized *Microcystis aeruginosa* PCC 7806 MysD with C-A pattern preferentially produces shinorine,^17^ we hypothesized that MysDs with X-A pattern (except for F-A) has higher specificity towards producing shinorine.

We expressed MysDs from *Microcystis aeruginosa* PCC 7806 (C-A), *Coleofasciculus chthonoplastes* PCC 7420 (C-A), *Cyanothece* sp. BG0011 (Y-A) and *Crocosphaera chwakensis* CCY0110 (Y-A) in SC.L26 and found out that all of them produce predominantly shinorine, suggesting a relationship between A in the second residue and shinorine product specificity with the exception of F-A (Figure 4B, Figure S19-S22). Similarly, we hypothesized that MysDs with X-S pattern (except for L-S) has higher specificity towards producing porphyra-334. We expressed MysDs from *Nostoc* sp. PA-18-2419 (Y-S) and *Nostoc* sp. NIES-4103 (Y-S), *Calothrix elsteri* CCALA 953 (A-S), *Calothrix* sp. PCC 6303 (A-S), *Chamaesiphon polymorphus* CCALA 037 (M-S) and *Chamaesiphon minutus* PCC 6605 (M-S) in SC.L26 and found out that all of them produce predominantly porphyra-334, suggesting a relationship between S in the second residue and porphyra-334 product specificity with the exception of L-S (Figure 4B, Figure S23-S28).

To validate that product specificity arises from key residue pattern rather than sequence similarity of whole enzymes, we aligned amino acid sequence of all characterized MysDs and constructed their phylogenetic tree (Figure S6). Sporadic distribution of MysDs with different product specificity throughout the tree was observed, suggesting independence of product specificity from sequence similarity.

Discovery of the relationship between pattern of key residues and product specificity enables us to screen for more specific shinorine and porphyra-334 producers with minimal side-product produced. Amongst all MysDs we characterized, MysD from *Microcystis aeruginosa* PCC 7806 (C-A) shows the highest shinorine-to-porphyra-334 ratio while MysD from *Nostoc* sp. PA-18-2419 (Y-S) shows the highest porphyra-334-to-shinorine ratio (Figure S4). These novel MysD orthologs hold significant potentials in the large-scale industrial synthesis of MAAs.

## Conclusion

In this study, we explored strategies to enable efficient and specific production of MAAs through pathway engineering in *S. cerevisiae*. Through introduction of xylose utilization pathway and multiple copies of MAA biosynthesis pathway genes and knock-down of endogenous competing pathways, we developed a yeast strain capable of producing MG, the precursor of all di-substituted MAAs. This yeast strain allows us to engineer shinorine and porphyra-334 production through expressing wild-type and mutant MysDs and explore MysD’s residue pattern-product specificity relationship. Using structure-guided mutagenesis approach, we corroborated on Kim et al., 2023’s result on the role of omega-loop in product specificity determination and took a step further, narrowing it down to the identity of only two residues. The residue pattern-product specificity correlation was supported by pattern-guided selection and characterization of MysD homologs. Hence, we can exploit the pattern of these two residues to predict, fine-tune and alter MysD product specificity. Our finding provides guidance for selecting uncharacterized MysD candidates for further screening and sheds light on the rational modifications on MysD that enables it to conjugate novel amino acid substrates.

## Materials and methods

### Protein Structure Prediction and Molecular Docking

3D structures of *Np*.MysD and *Nl*.MysD were predicted using ColabFold online server.^19^ The predicted structures were then aligned with the crystallographic structure of *Escherichia coli* d-alanine-d-alanine ligase (PDB ID: 2DLN). Ligand model was constructed and visualized using ChemOffice Professional, Version 20.0 PerkinElmer Informatics, 2020. Structural alignment and visualization of proteins were performed using The PyMOL Molecular Graphics System, Version 2.0 Schrödinger, LLC. AutoDockTools 1.5.7 (ADT) was used to pre-process enzyme and ligand files and Autodock Vina 1.2.0 was used to dock ligand into the active site of the enzymes. ^20–22^ The molecular conformations with the highest affinity score (lowest kcal/mol) were used for the subsequent analysis.

### Multi-sequence Alignment and the Construction of Phylogenetic Tree

The amino acid sequence of *Np*.MysD and *Nl*.MysD were acquired from the NCBI Nucleotide Database (GenBank: ACC83902.1, AP018223.1).^23^ The sequences were imported into MEGA v11.0 and aligned using the MUSCLE algorithm with default parameters.^24^

The amino acid sequences of the 20 cyanobacterial mysD orthologs were either acquired from the OrthoDB v11 Database or the NCBI Protein Database.^23,25^ The sequences of the Omega-Loops were extracted from the original sequence and aligned using the MUSCLE algorithm with default parameters in MEGA v11.0. To construct the Maximum-Likelihood (ML) tree, the software IQ-TREE was used and the JTTDC model was applied, with additional parameters +I and +G (Invariable site plus discrete Gamma model) and 1000-UltraFast Bootstrap Approximation (UFBoot)^26,27^. Other iq-tree parameters were set as default. The exported phylogenetic tree was then processed and annotated using iTOL v6.35.^28^

### Engineering MAA production yeast strains

The exogenous genes used in this study (*xyl1-3, Np*.*mysA-D, Nl*.*mysD, 20 new characterized MysDs*) were synthesized from the BGI Genomics. The sequences were flanked with BsaI restriction sites on both termini and assembled onto the intermediate plasmid (type-3).

Plasmids link-021, link-022, link-024, link-027 and (see Table 2) contain various promoter and terminator elements, with BsaI sites compatible to the intermediate plasmids. The expression vectors were constructed using the NEBridge® Golden Gate Assembly Kit (BsaI-HF® v2). The knockout of *Tal1* and *Nqm1* and the genomic insertion of *xyl1-3* and *mysA-C in S*.*cerevisiae* were implemented using CRISPR-Cas9.^29^ For genomic insertion, the corresponding DNA fragment and the pCRCT plasmids^30^ (Table 2) were chemically transformed into *S*.*cerevisiae*, followed by verification procedures including colony PCR and sequencing.

### Point-mutations and omega-loop switching

Primers were designed for site-directed mutagenesis (Table S4). Pairs of point-mutation primers containing the identical 5’ overhangs introduce homology arms during PCR. The linear DNA fragments were then circularized into plasmids using the Gibson Assembly® Cloning Kit from New England Biolabs. For omega-Loop exchanges, two rounds of PCR were applied. The first round generates linear DNA fragments with partially-exchanged omega-loop sequences and no homology arms, while the second round generates completely-exchanged omega-loop sequences and the homology arms used for Gibson Assembly® (Table S4). Success of mutation and loop change were determined through Sanger DNA sequencing.

### Assembly of *mysD* genes from other cyanobacteria strains

20 different naturally found mysD genes on NCBI were synthesized by BGI Genomics and inserted on links-024s vector. Synthesized plasmids were transformed into SC.L26 for plasmid expression and production of porphyra-334 and shinorine.

### Growth of *S. cerevisiae* strains and the extraction method

Strains used in this study are listed in Table-1. *S. cerevisiae* strains were cultured in SC culture medium without uracil (1.7 g/L yeast nitrogen base without amino acids, 5 g/L Ammonium sulfate and 1.29 g/L DO Supplement -Ura) containing total of 12g/L of glucose and 8g/L xylose. For extraction of MAA samples, suspension of the *S*.*cerevisiae* cultures were harvested and freeze-thawed twice after 48 hours of fermentation. After centrifugation, the supernatant was retained and filtered with 0.22 um filtration membrane. The processed samples were stored at -20°C before the LC-HRMS analysis.

**Table 1.** Strains used in this study.

| Strain | Genotype | Reference |
| --- | --- | --- |
| CEN.PK2 | MATa ura3-52 trp1-289 leu2-3112 his3Δ1 MAL2-8<sup>c</sup> SUC2 | EUROSCARF |
| SC.L1 | CEN.PK2 Tal1Δ | This study |
| SC.L2 | SC.L1 His3::P<sub>TDH3</sub> - *XYL1* -T<sub>TDH1</sub>-P<sub>PGK1</sub>-*XYL2*-T<sub>CYC1</sub>-P<sub>TEF2</sub>-*XYL3*-T<sub>SSA1</sub> | This study |
| SC.L3 | SC.L2 308::P<sub>TDH3</sub>-*np.mysA*-T<sub>TDH1</sub>-P<sub>PGK1</sub>-*np.mysB*-T<sub>PGK1</sub> | This study |
| SC.L6 | SC.L3 Nqm1::P<sub>TDH3</sub>-*np.mysB*-T<sub>TDH1</sub>-P<sub>PGK1</sub>-*np.mysA*-T<sub>PGK1</sub> | This study |
| SC.L26 | SC.L6 YP::P<sub>TDH3</sub>-*np.mysC*-T<sub>TDH1</sub> | This study |

**Table 2.** Plasmids used in this study.

| Plasmid | Description | Reference |
| --- | --- | --- |
| links021 | 2U-URA3-LS-P<sub>TDH3</sub>-Bsal-T<sub>TDH1</sub>-R1 | This study |
| links024 | 2U-URA3-L1-P<sub>PGK1</sub>-Bsal-T<sub>PGK1</sub>-R2 | This study |
| links027 | 2U-URA3-L3-P<sub>TEF2</sub>-Bsal-T<sub>SSA1</sub>-RE | This study |
| links022 | 2U -URA3-P<sub>PGK1</sub>-Bsal-T<sub>CYC1</sub>-R2 | This study |
| type3-XYL1 | M13F-Bsal-XYL1-Bsal-M13R | This study |
| type3-XYL2 | M13F-Bsal-XYL2-Bsal-M13R | This study |
| type3-XYL3 | M13F-Bsal-XYL3-Bsal-M13R | This study |
| type3-Np.mysA | M13F-Bsal-Np.mysA-Bsal-M13R | This study |
| type3-Np.mysB | M13F-Bsal-Np.mysB-Bsal-M13R | This study |
| type3-Np.mysC | M13F-Bsal-Np.mysC-Bsal-M13R | This study |
| type3-Np.mysD | M13F-Bsal-Np.mysD-Bsal-M13R | This study |
| type3-Nl.mysD | M13F-Bsal-Nl.mysD-Bsal-M13R | This study |
| type9K-XYL1-XYL2-XYL3 | P<sub>TDH3</sub>-XYL1-T<sub>TDH1</sub>-P<sub>PGK1</sub>-XYL2-T<sub>CYC1</sub>-P<sub>TEF2</sub>-XYL3-T<sub>SSA1</sub> | This study |
| pCRCT-TAL1 | ura3, P<sub>SNR52</sub>-TAL1-T<sub>sup4</sub>, P<sub>TEF1</sub>-CAS9-T<sub>RPR1</sub> flanked by TAL1 upstream and downstream | This study |
| pCRCT-HIS3 | ura3, P<sub>SNR52</sub>-HIS3-T<sub>SUP4</sub>, P<sub>TEF1</sub>-CAS9-T<sub>RPR1</sub> | This study |
| pCRCT-308a | ura3, P<sub>SNR52</sub>-308a-T<sub>SUP4</sub>, P<sub>TEF1</sub>-CAS9-T<sub>RPR1</sub> | This study |
| pCRCT-YP | ura3, P<sub>SNR52</sub>-YPRCd15c-T<sub>SUP4</sub>, P<sub>TEF1</sub>-CAS9-T<sub>RPR1</sub> | This study |
| pCRCT-Nqm1 | ura3, P<sub>SNR52</sub>-Nqm1-T<sub>SUP4</sub>, P<sub>TEF1</sub>-CAS9-T<sub>RPR1</sub> | This study |

### LC-HRMS analytical methods

LC-HRMS analysis was performed on a UHPLC Vanquish flex system, equipped with a variable wavelength detector (Thermo Fisher Scientific, Bremen, Germany), coupled to a Q-Exactive Plus mass spectrometer (Thermo Fisher Scientific, Bremen, Germany) and operated in the positive (ESI+) and negative (ESI-) electrospray ionization modes (one run for both modes). The system was controlled by Xcalibur 4.2 (Thermo Fisher Scientific). The HESI (heated electrospray ionization) source used, spray voltages of 3.0 kV for ESI+ and -2.8 kV for ESI-, capillary temperature of 350 °C, heater temperature of 350 °C, sheath gas flow of 50 arbitrary units (AU), auxiliary gas flow of 15 AU, sweep gas flow of 2 AU, and S-Lens RF level of 50%. During the full-scan acquisition, which ranged from m/z 100 to 800, the instrument operated at 70,000 resolution at m/z = 200, with an automatic gain control (AGC) target of 3 × 106 and a maximum injection time (MIT) of 100 ms. For MS2 analysis, the isolation window was set at 4.0 m/z, the instrument was operated at 17,500 resolution at m/z = 200, with an AGC target of 1 × 105, MIT of 50 ms, and stepped NCE of 30, 40, and 50 eV with N2 as collision gas.

Chromatographic separation was carried out with a UPLC HSS T3 column (2.5 µm, 100 mm ×2.1 mm), at a temperature of 25 °C. The mobile phases consisted of 0.1% formic acid in water (A) and acetonitrile (B). The elution gradient was as follows: 0-5 min, 1% B; 5-7 min, 1-30% B; 7-9 min, 30-99% B; 9-10 min, 99% B; 10-10.1 min, 99-1% B; 10.1-15 min, 1% B, at aflow rate of 0.2 mL/min. The injection volume was 1 µL. UV absorption was set at a fixed wavelength of 334 nm.

### Peak area quantification

Shinorine and porphyra-334 standard samples were acquired from Beijing Yiming Fuxing Biological Technology Co., Ltd. Concentrations of shinorine and porphyra-334 were calculated by comparing peak areas produced by each sample with the peak areas of 0.1 mg/mL standards. Response factor normalization was applied to calculated peak areas if shinorine and porphyra-334 showed difference in response factors during LC-MS.

## Acknowledgement

The authors are grateful to E Li from the Multi-Omics Mass Spectrometry Core of Shenzhen Bay Laboratory for conducting the mass spectrometry experiment and analysis of the MS data. We thank Jun Yu for providing advice to experimentation. We thank Xinxin Lu for acquisition of shinorine and porphyra-334 standards. HS is supported by the BBSRC Norwich Research Park Doctoral Training Partnership Scholarship (BB/T008717/1 Project No. 2578291).

## Author contributions

XZ, HS and BW conceptualized this study. XZ, PX, ACC, YJ, SH and XM performed investigation. XZ, PX, HS and BW performed formal analysis and visualization and contributed to writing and editing. HS and BW were responsible for supervision. BW was responsible for funding acquisition.

