## Supplemental Figures for "Decoding Specificity of Cyanobacterial MysDs in Mycosporine-Like Amino Acid Biosynthesis through Heterologous Expression in *Saccharomyces cerevisiae*": MAA_Supplementary Materials.docx

**Glossary.** Abbreviations and full names of the metabolites involved in this study.

| **Abbreviation** | **Full Name** |
| --- | --- |
| G6P | Glucose 6-phosphate |
| F6P | Fructose 6-phosphate |
| F1,6P | Fructose 1,6-bisphosphate |
| G3P | Glyceraldehyde 3-phosphate |
| DHAP | Dihydroxyacetone phosphate |
| 1,3PG | 1,3-Bisphosphoglycerate |
| 6PGL | 6-Phosphogluconolactone |
| 6PG | 6-Phosphogluconate |
| Ru5P | Ribulose 5-phosphate |
| R5P | Ribose 5-phosphate |
| X5P | Xylulose 5-phosphate |
| S7P | Sedoheptulose 7-phosphate |
| E4P | Erythrose 4-phosphate |
| DDG | Desmethyl 4-deoxygadusol |
| 4-DG | 4-Deoxygadusol |
| MG | Mycosporine-glycine |

**Table S1.** Genes used in metabolic engineering.

| ***gene name*** | **ACCESSION** | **organism name** |
| --- | --- | --- |
| *XYL1* | XP_001385181 | *Scheffersomyces stipitis* CBS 6054 |
| *XYL2* | XP_001386982 | *Scheffersomyces stipitis* CBS 6055 |
| *XYL3* | XP_001387325 | *Scheffersomyces stipitis* CBS 6056 |
| *mysA(DDG)* | WP_012411849 | *Nostoc punctiforme* |
| *mysB(O-MT)* | WP_012411848 | *Nostoc punctiforme* |
| *mysC(AGL)* | WP_012411847 | *Nostoc punctiforme* |
| *Np.mysD* | WP_012411846 | *Nostoc punctiforme* |
| *Nl.mysD* | BAY79928 | *Nostoc linckia* NIES-25 |

**Table S2.** sgRNA Sequences.

| **Target site** | **Guide Sequence** |
| --- | --- |
| TAL1 | ACTGTCGTTGTTGCCGACAC |
| HIS3 | TGCCTCGCAGACAATCAACG |
| 308a | CACTTGTCAAACAGAATATA |
| Nqm1-1 | GATCAAGATAGCTTCTACGT |
| YPRCd15c | AATCCGAACAACAGAGCATA |

**Table S3.** Novel *mysDs* from 20 cyanobacteria strains.

| ***gene name*** | **ACCESSION** | **organism name** | **Source** |
| --- | --- | --- | --- |
| *CA-267872*  *FS-450202*  *LA-76335* | WKX64033.1  BBB38339.1  WP_114083456 | *Microcystis aeruginosa PCC 7806*  *Nostoc commune KU002*  *Nostoc sp.* ATCC 53789 | NCBI Proteins  NCBI Proteins  OrthoDB v11 |
| *LA-46234* | WP_015081019 | *Anabaena sp.* 90 | OrthoDB v11 |
| *FS-1932621* | WP_086687004 | *Nostoc sp.* T09 | OrthoDB v11 |
| *FS-1628751* | PHK43867 | *Nostoc linckia* z16 | OrthoDB v11 |
| *FS-446679* | WP_118166108 | *Nostoc sphaeroides* | OrthoDB v11 |
| *FA-102232* | WP_006527735 | *Gloeocapsa sp.* PCC 73106 | OrthoDB v11 |
| *CA-118168* | WP_006100251 | *Coleofasciculus chthonoplastes* PCC7420 | OrthoDB v11 |
| *AS-1170562* | WP_015199129 | *Calothrix sp.* PCC 6303 | OrthoDB v11 |
| *AS-987040* | WP_095720873 | *Calothrix elsteri* CCALA 953 | OrthoDB v11 |
| *YS-2575443* | WP_138500196 | *Nostoc sp.* PA-18-2419 | OrthoDB v11 |
| *YS-2005458* | BAZ51206 | *Nostoc sp.* NIES-4103 | OrthoDB v11 |
| *YA-2082950* | WP_107668801 | *Cyanothece sp.* BG0011 | OrthoDB v11 |
| *YA-391612* | WP_008274182 | *Crocosphaera chwakensis* CCY0110 | OrthoDB v11 |
| *MS-2107692* | PSB40329 | *Chamaesiphon polymorphus* CCALA 037 | OrthoDB v11 |
| *MS-1173020* | WP_015160004 | *Chamaesiphon minutus* PCC 6605 | OrthoDB v11 |
| *LS-179408* | WP_015177382 | *Oscillatoria nigro-viridis* PCC 7112 | OrthoDB v11 |
| *LA-457944* | WP_069074327 | *Nostoc sp.* KVJ20 | OrthoDB v11 |
| *LA-163908* | WP_016949469 | *Anabaena sp.* PCC 7108 | OrthoDB v11 |

**Table S4.** Primers for omega-loop exchanging and point-mutagenesis.

| **Template** | **Primer Pair** | **Sequence (5’-3’)** | **Amplicon** |
| --- | --- | --- | --- |
| links024-*Np.mysD* | *Np.*Loop-A-F | GTTTCGCTGCCAAGGGTAACAACAAATCTTGGATTTTAGACCCTAACGACC | Fragment-1. |
|  | *Np.*Loop-A-R | CATCAGTTTTCTTCAACTTGTCTGCGTAGGTACGGATAGGTTTATCGTGTGG |  |
| Fragment-1 | *Np.*Loop-B-F | AAAACTGATGACGGTTCTTTAGGTTTCGCTGCCAAGGGTA | Fragment-1 with two homology arms. Became links024-*Np.mysD-Nl.*Loop after Gibson assembly. |
|  | *Np.*Loop-B-R | AGCGAAACCTAAAGAACCGTCATCAGTTTTCTTCAACTTGTCTG |  |
| links024-*Np.mysD* | *Np*.LmutF-F | GCGACTTGCATTTCACTGCTAAAGATAATATCAAGGCTTGG | Fragment-2. Became links024-*Np.mysD*-L249F after Gibson assembly. |
|  | *Np.*LmutF-R | TATTATCTTTAGCAGTGAAATGCAAGTCGCCATCGTCA |  |
| links024-*Np.mysD* | *Np.*AmutS-F | AGATAATATCAAGTCTTGGATTTTAGACCCTAACGACC | Fragment-3. Became links024-*Np.mysD*-A257S after Gibson assembly. |
|  | *Np.*AmutS-A-R | GGGTCTAAAATCCAAGACTTGATATTATCTTTAGCAGTCAAATGCAAG |  |
| links024-*Np.mysD*-L249F | *Np.*AmutS-F | AGATAATATCAAGTCTTGGATTTTAGACCCTAACGACC | Fragment-4. Became links024-*Np.mysD*-L249F-A257S after Gibson assembly. |
|  | *Np.*AmutS-B-R | GGGTCTAAAATCCAAGACTTGATATTATCTTTAGCAGTGAAATGCAAG |  |
| links024-*Nl.mysD* | *Nl.*Loop-A-F | GCATTTGACTGCTAAAGATAATATCAAGGCTTGGATTTTGGACCCAAACGA | Fragment-5. |
|  | *Nl.*Loop-A-R | CGTCAGTTTGTTGGAGTTTATCAGCATAGTTTCTGATTGGTTTTTCTTGAGAGTCG |  |
| Fragment-5 | *Nl.*Loop-B-F | CAAACTGACGATGGCGACTTGCATTTGACTGCTAAAGATAATATCAAGG | Fragment-5 with two homology arms. Became links024-*Nl.mysD-Np.*Loop after Gibson assembly. |
|  | *Nl.*Loop-B-R | AGTCAAATGCAAGTCGCCATCGTCAGTTTGTTGGAGTTTATCAG |  |
| links024-*Nl.mysD* | *Nl.*FmutL-F | GTTCTTTAGGTTTAGCTGCCAAGGGTAACAACAA | Fragment-6. Became links024-*Nl.mysD*-F247L after Gibson assembly. |
|  | *Nl.*FmutL-R | CCCTTGGCAGCTAAACCTAAAGAACCGTCATCAGTT |  |
| links024-*Nl.mysD* | *Nl.*SmutA-F | AAGGGTAACAACAAAGCTTGGATTTTGGACCCAAACGA | Fragment-7. Became links024- *Nl.mysD*-S255A after Gibson assembly. |
|  | *Nl.*SmutA-R | GTCCAAAATCCAAGCTTTGTTGTTACCCTTGGCAGC |  |
| links024-*Nl.mysD*-F247L | *Nl.*SmutA-F | AAGGGTAACAACAAAGCTTGGATTTTGGACCCAAACGA | Fragment 8. Became links024-*Nl.mysD*-F247L-S255A after Gibson assembly. |
|  | *Nl.*SmutA-R | GTCCAAAATCCAAGCTTTGTTGTTACCCTTGGCAGC |  |

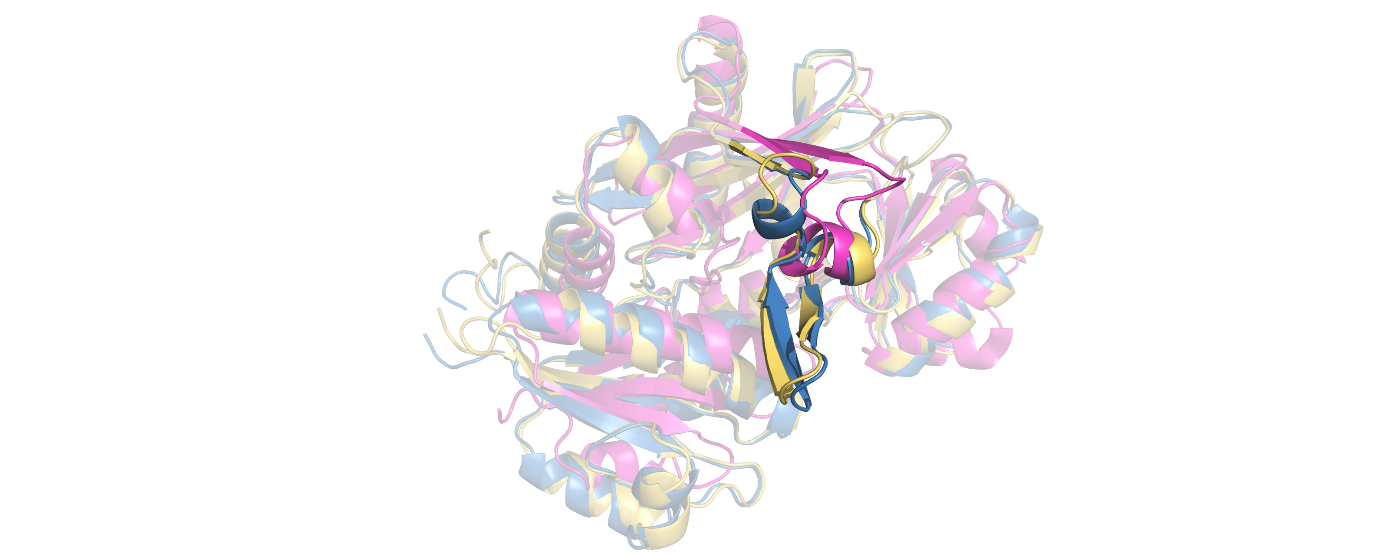

**Figure S1.** Structural alignment of the predicted structural model of *Nl.*MysD (blue) and the predicted structural model of *Np.*MysD (yellow) with the crystal structure (magenta) of Escherichia coli d-alanine-d-alanine ligase (2DLN). The omega-loops are highlighted with opacity while the rest of the structures are transparent. Water and ligand molecules from the original PDB file of 2DLN are removed.

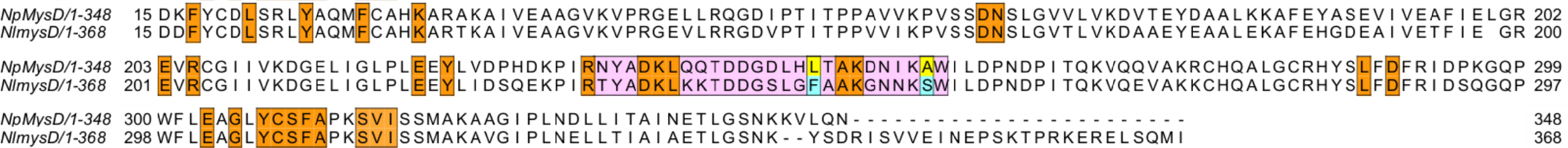

**Figure S2.** The complete version of the aa sequence alignment between *Np.*MysD and *Nl.*MysD. The identical residues within 5 angstroms from the docked reaction intermediate (P-MG-Ser) are colored orange. The two critical aa residues are highlighted with yellow (*Np.*MysD) or cyan (*Nl.*MysD). The 26-aa long fragment of the omega-loop (pink) contains all the variable aa residues within 8 angstroms.

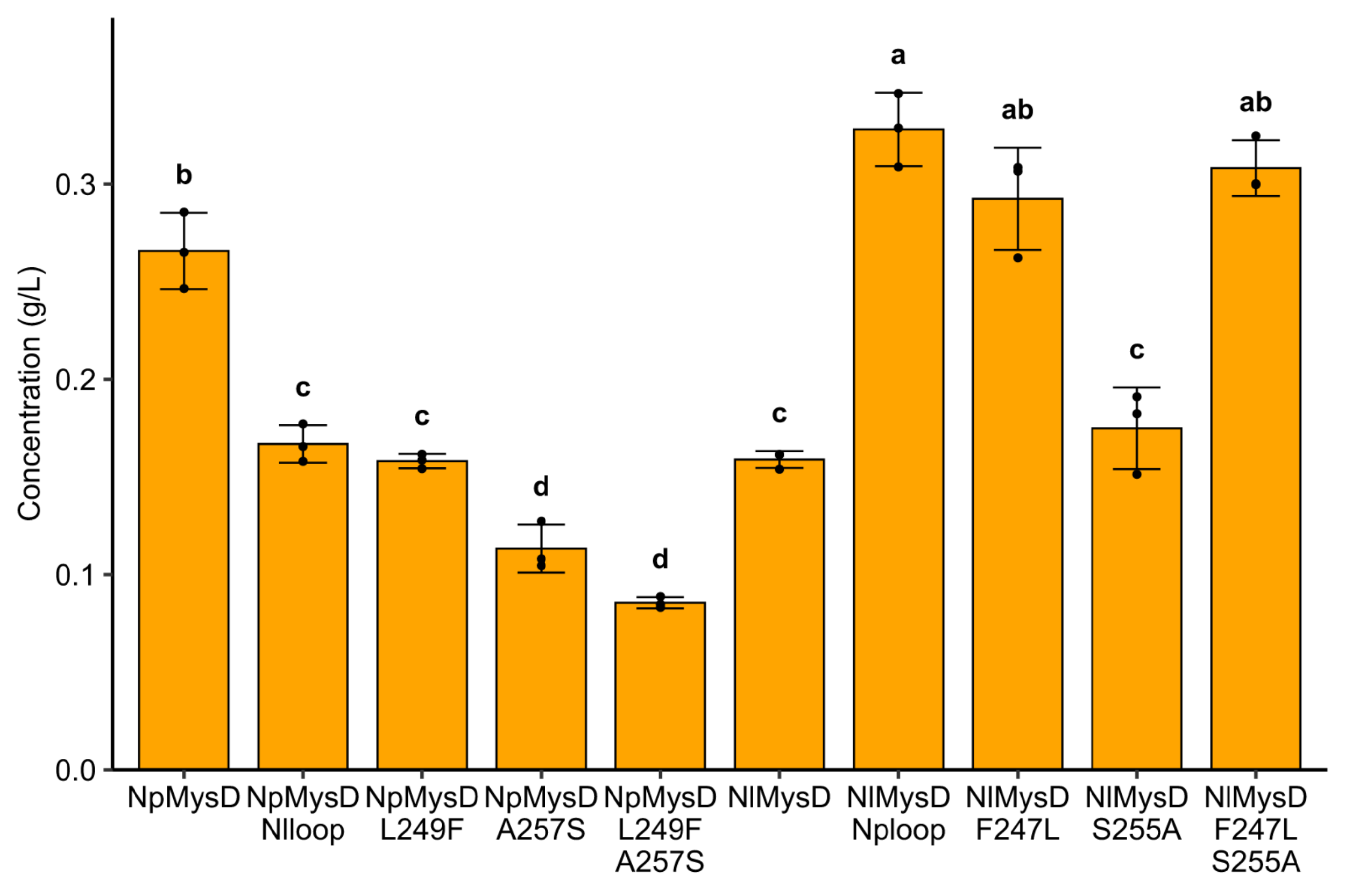

**Figure S3.** Quantification of total enzyme activity by comparing the sum of shinorine and porphyra-334 peak area amongst these yeast strains. Heights of the bars represent their means and the error bars represent their standard deviations (n=3 biological replicates). One-way ANOVA was performed to compare the total amount of shinorine and porphyra-334 amongst different samples and two samples that do not share the same letter are significantly different from each other (p<0.05).

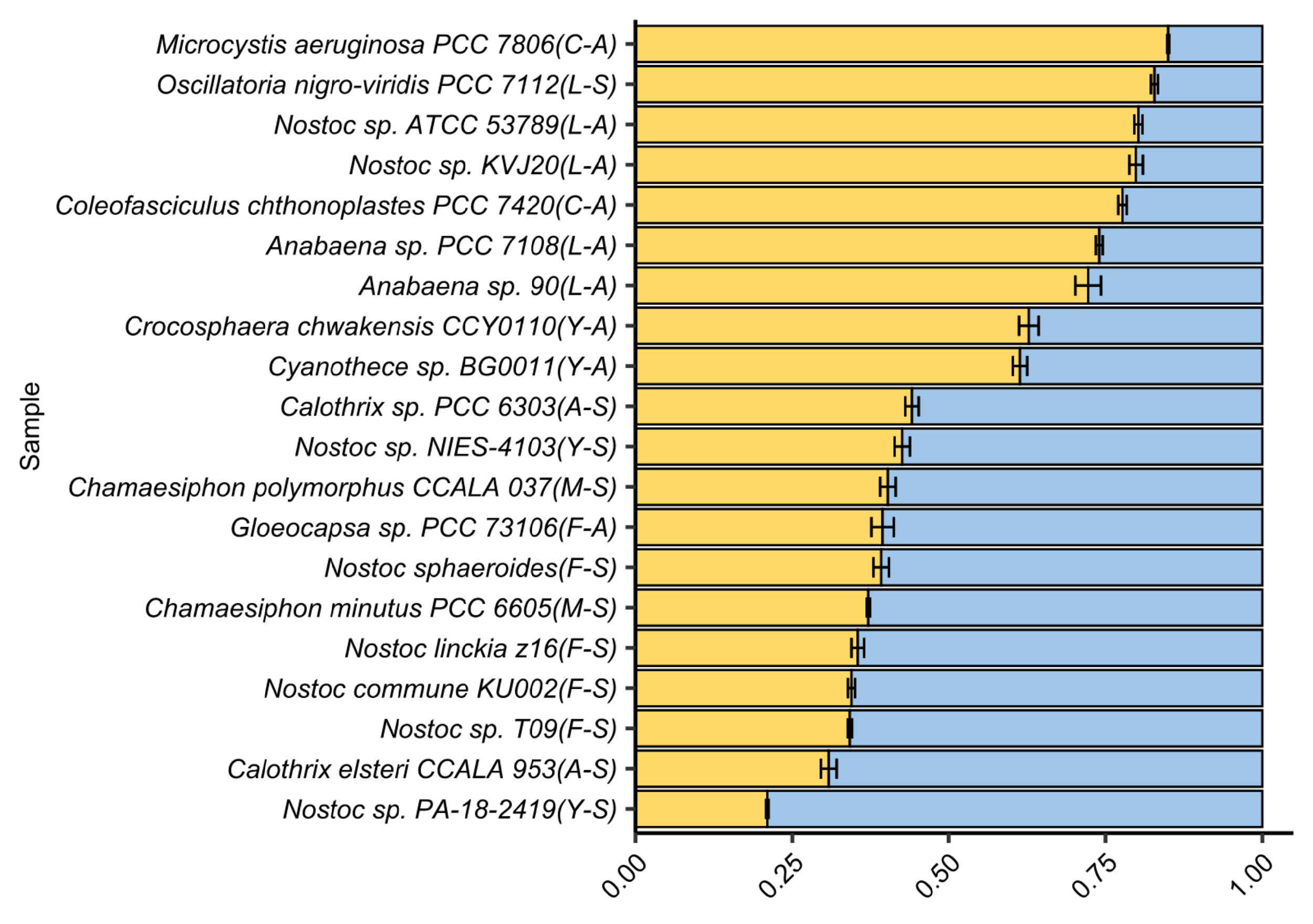

**Figure S4.** Shinorine (yellow) : porphyra-334 (blue) ratio from LC-HRMS peak area analysis by yeast strains expressing selected MysDs. Heights of the bars represent their means and the error bars represent their standard deviations (n=3 biological replicates).

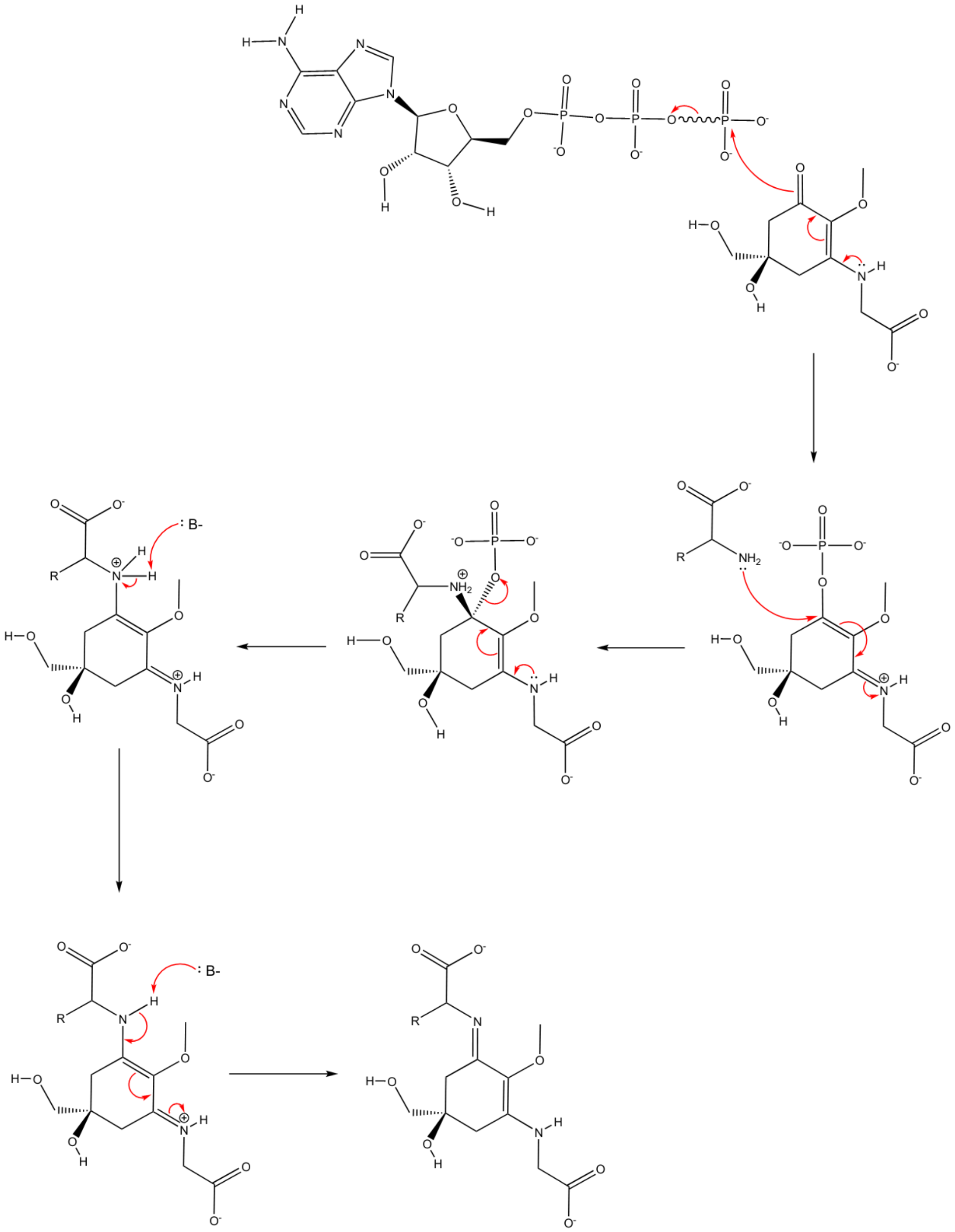

**Figure S5.** Proposed step-wise mechanism of the reaction catalyzed by MysDs. The process is initiated by the transfer of the terminal phosphate group from ATP to the C1 position of MG. The amino acid substrate is then conjugated with MG via a nucleophilic attack on C1. Through series of molecular rearrangements and deprotonations, the reaction is completed with the removal of the phosphate group.

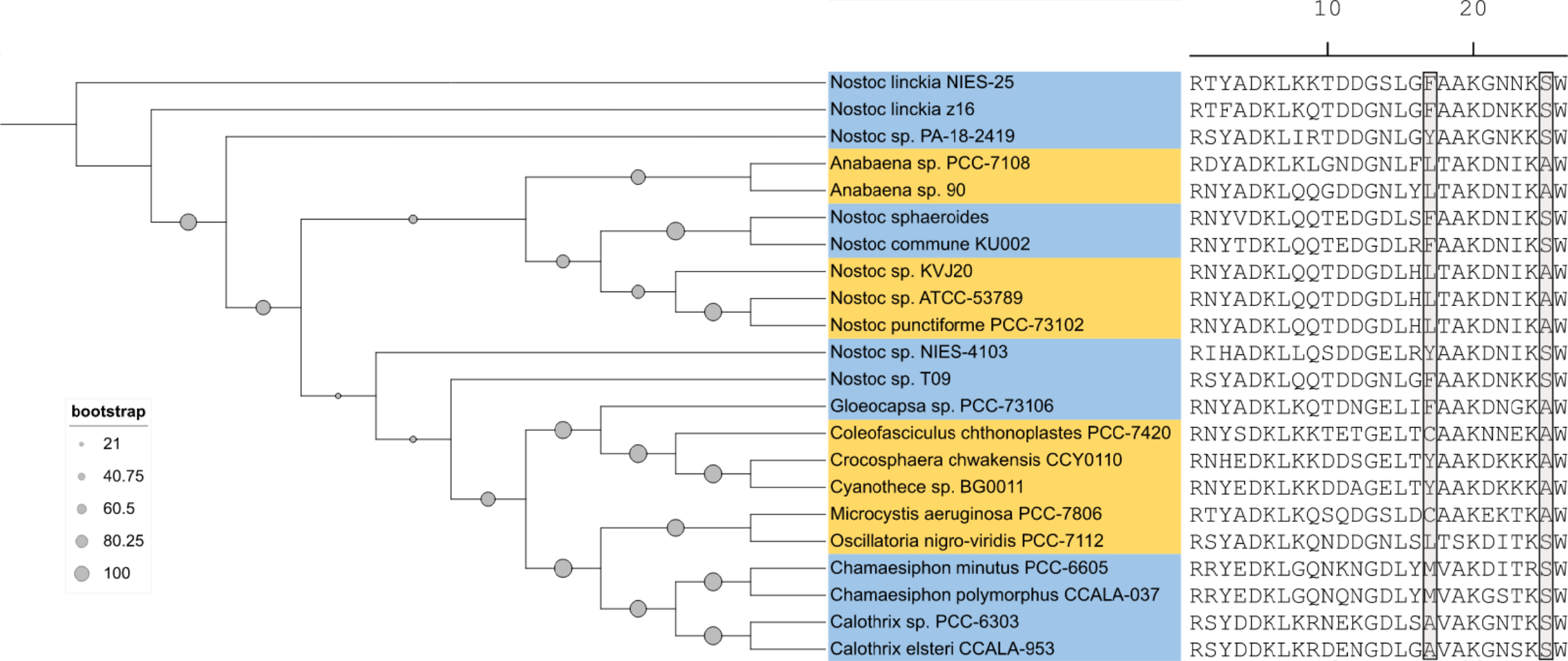

**Figure S6.** The Maximum-Likelihood (ML) phylogenetic tree of the 20 MysD orthologs. The tree was constructed using whole-sequence alignment and the 26-aa omega-loop fragments are shown. The bootstrap value of each node in the tree is demonstrated. The colors superimposed on the name of the strains indicate their product-specificity analyzed with LC-HRMS peak area. Blue indicates porphyra-334 preference, while yellow indicates shinorine preference. The two critical residues are labeled with grey boxes.

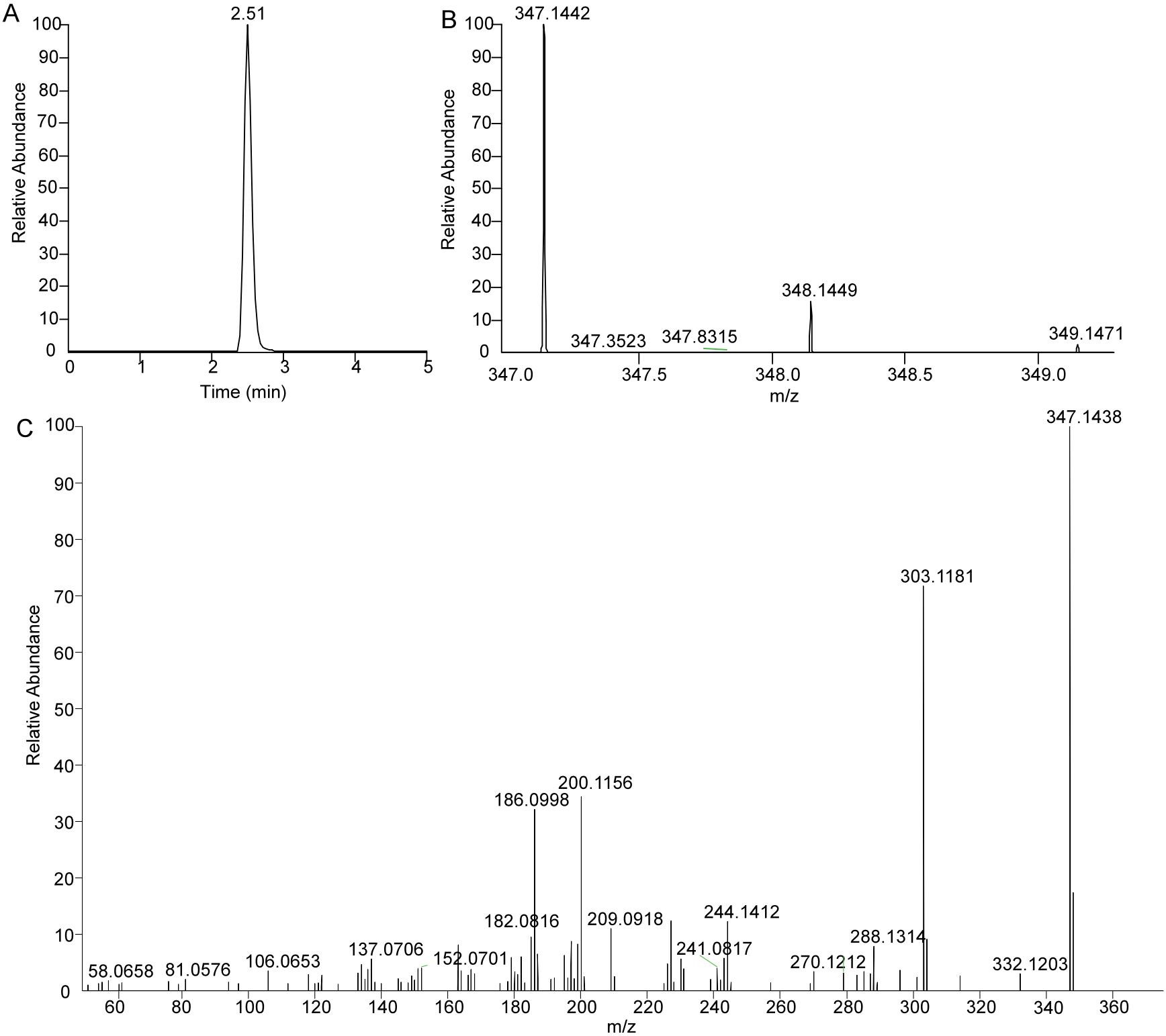

**Figure S7.** The extracted chromatogram peak, MS^1^ and MS/MS spectra of porphyra-334 for the characterization of the 20 MysD orthologs (A-C).

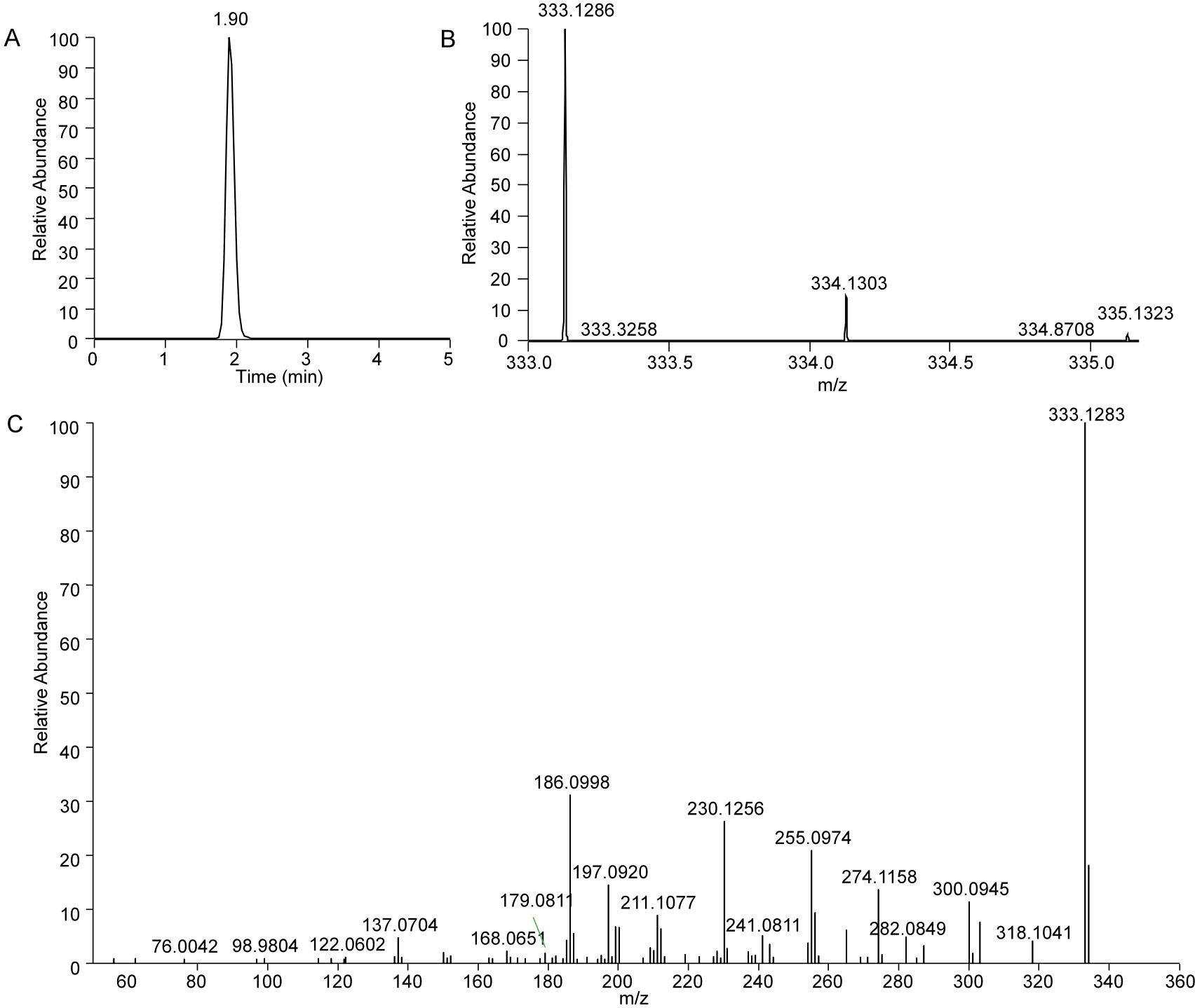

**Figure S8.** The extracted chromatogram peak, MS^1^ and MS/MS spectra of shinorine for the characterization of the 20 MysD orthologs (A-C).

**Figure S9-S28.** Chromatogram peaks of shinorine and porphyra-334 in the 20 MysD orthologs.

**S9. Anabaena sp. 90(L-A)**

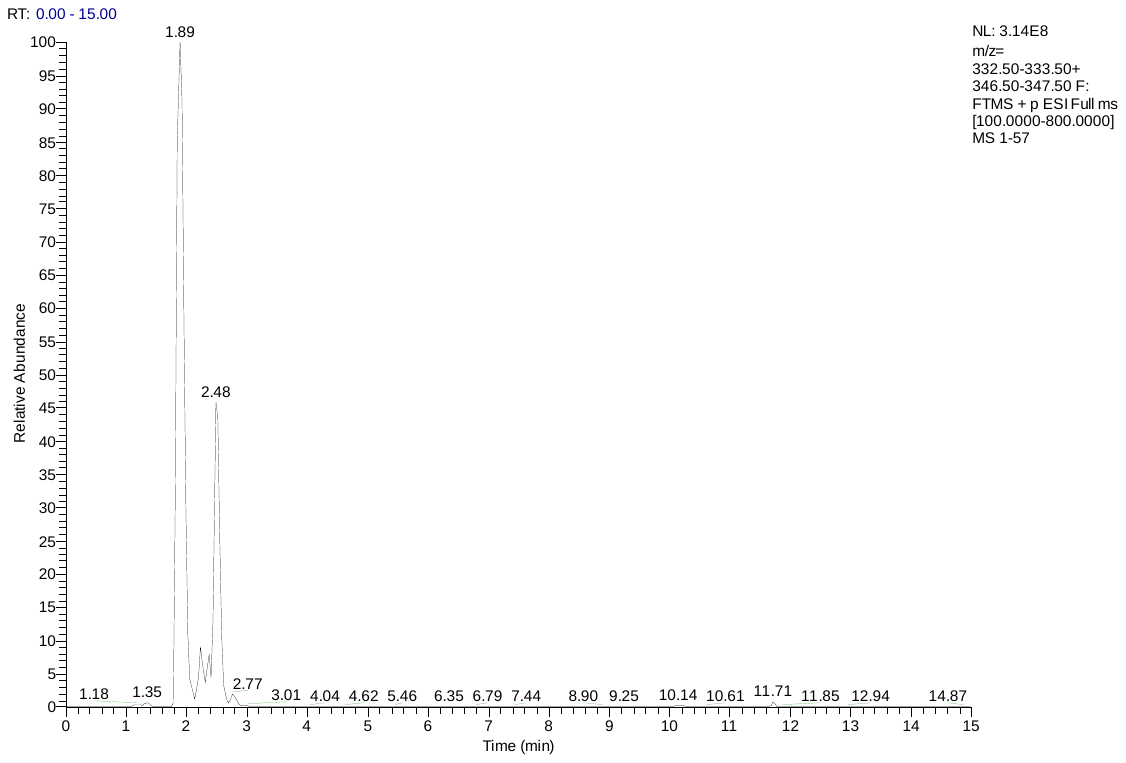

**S10. Anabaena sp. PCC 7108(L-A)**

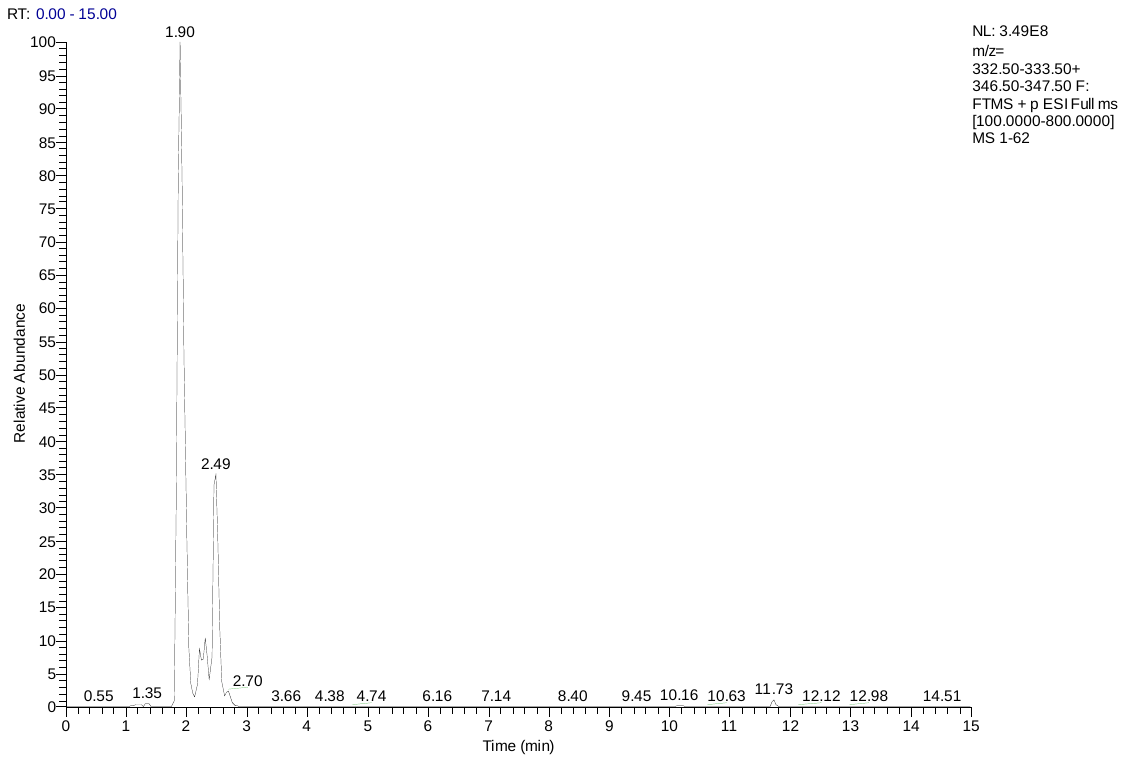

**S11. Nostoc sp. KVJ20(L-A)**

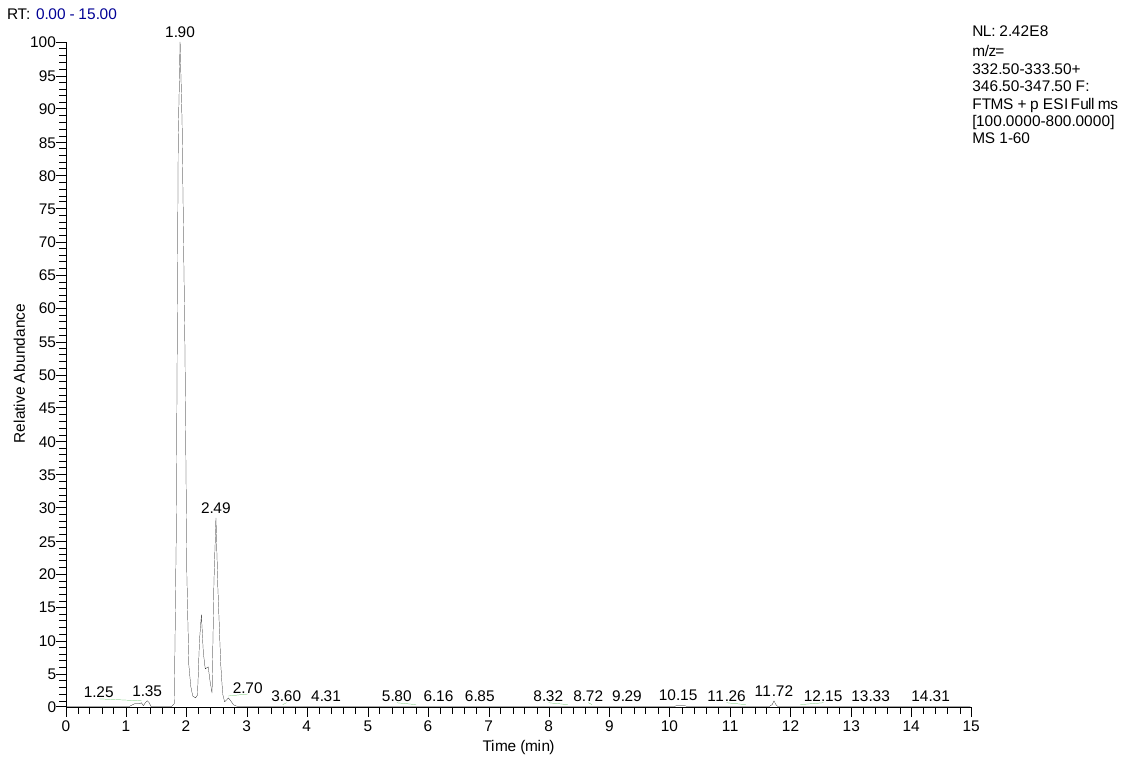

**S12. Nostoc sp. ATCC 53789(L-A)**

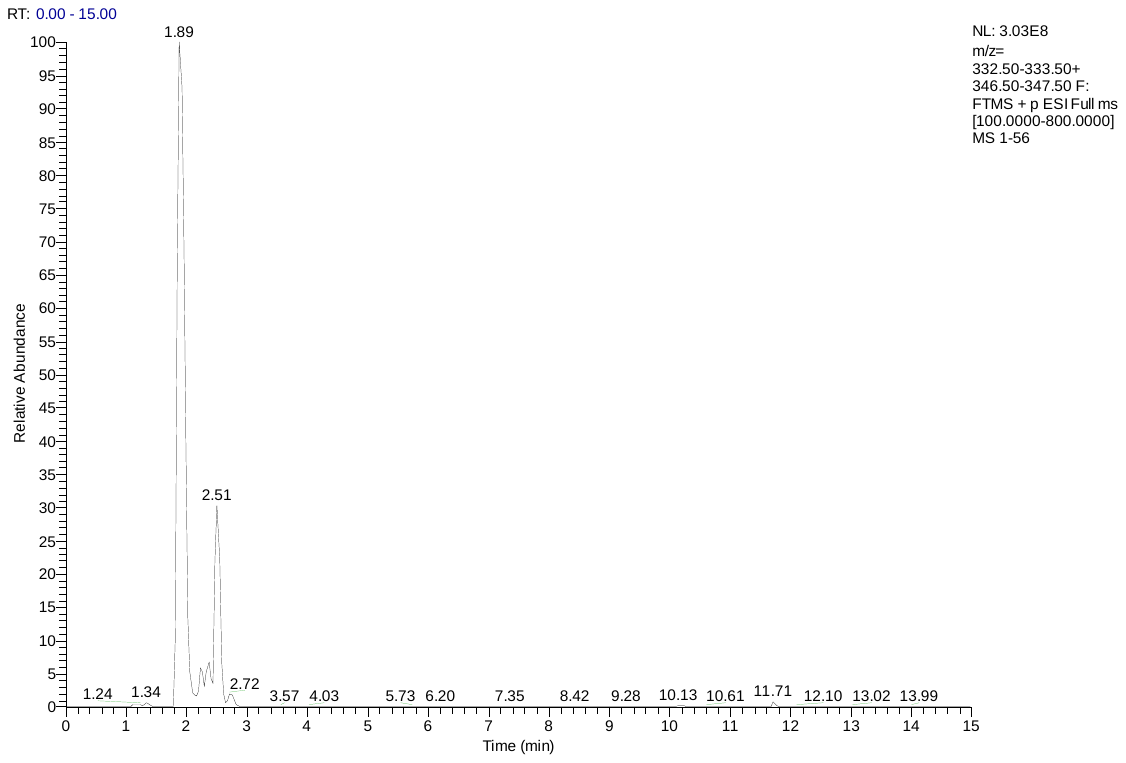

**S13. Nostoc linckia z16(F-S)**

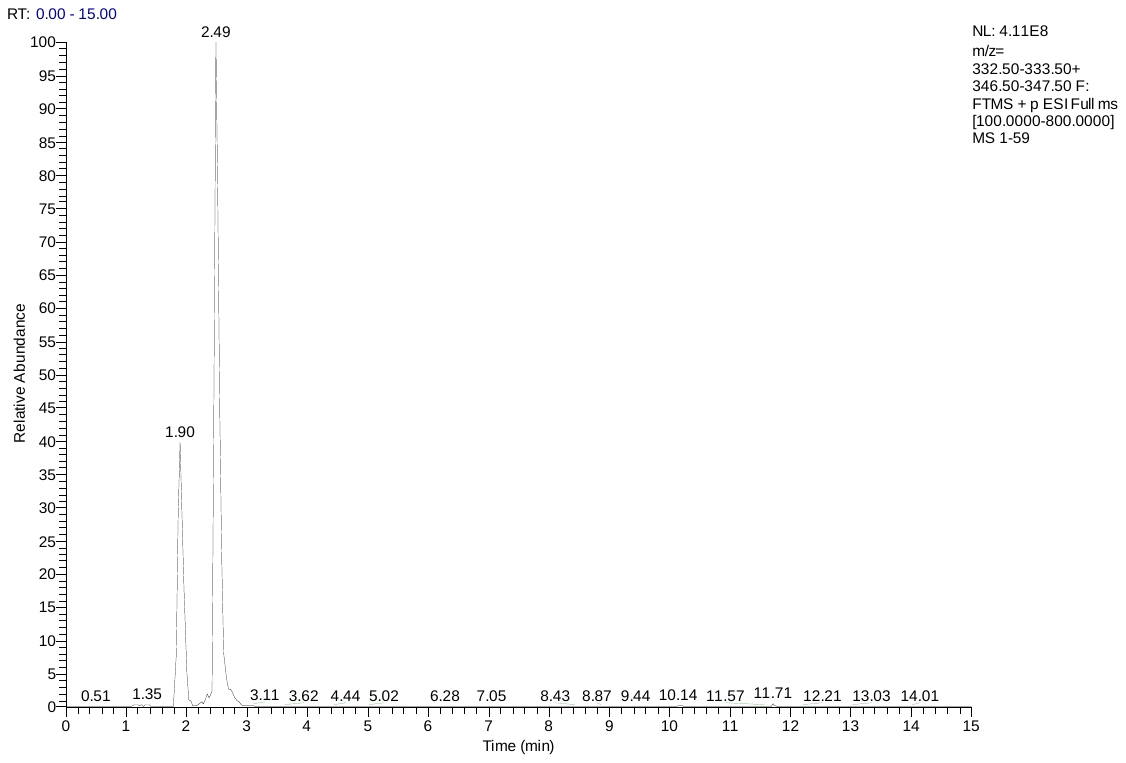

**S14. Nostoc sp. T09(F-S)**

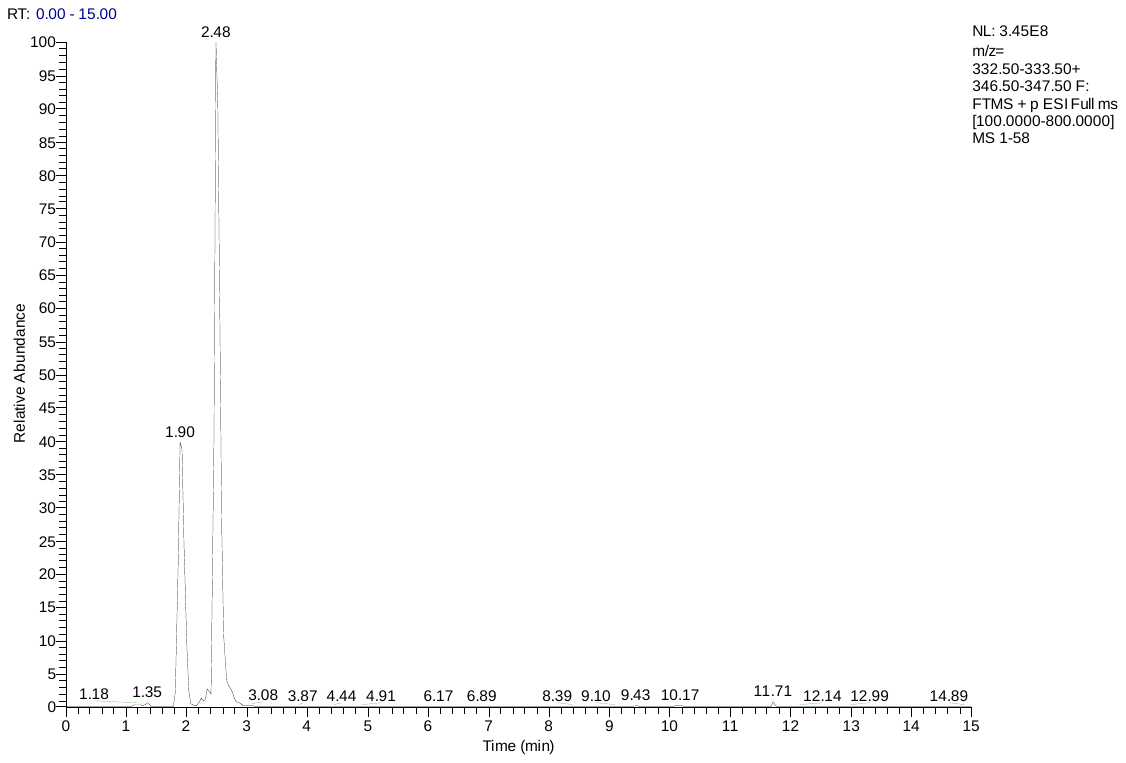

**S15. Nostoc commune KU002(F-S)**

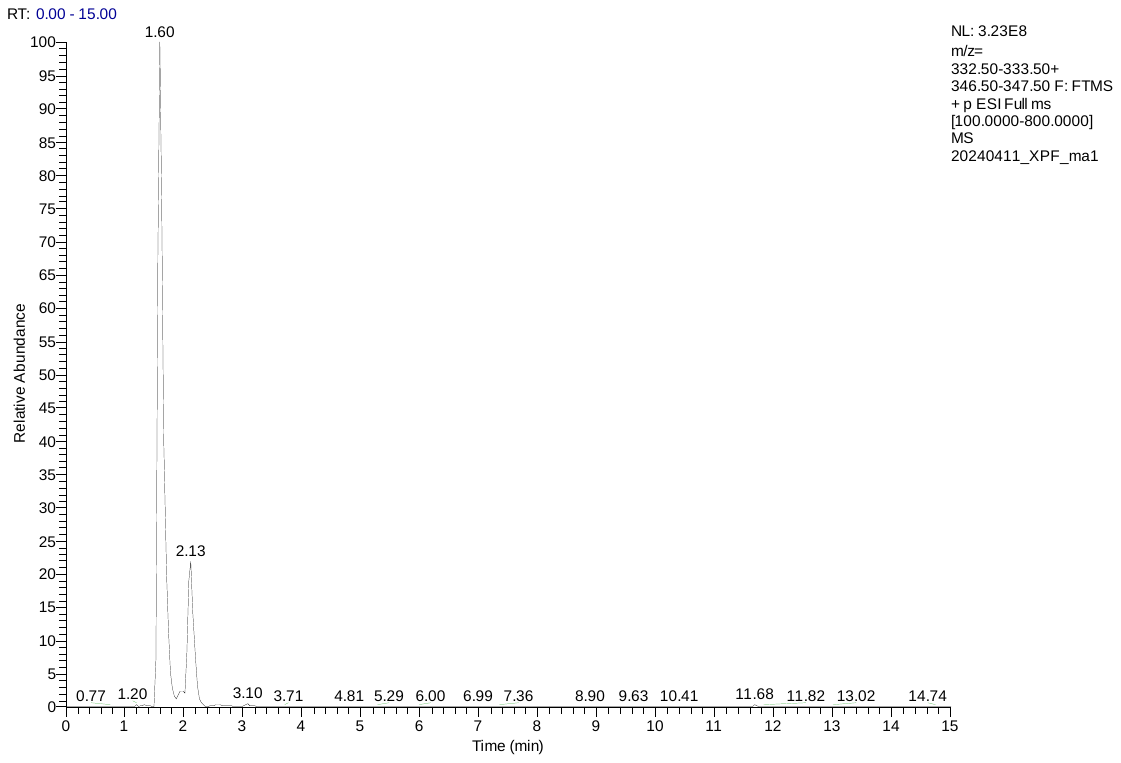

**S16. Nostoc sphaeroides(F-S)**

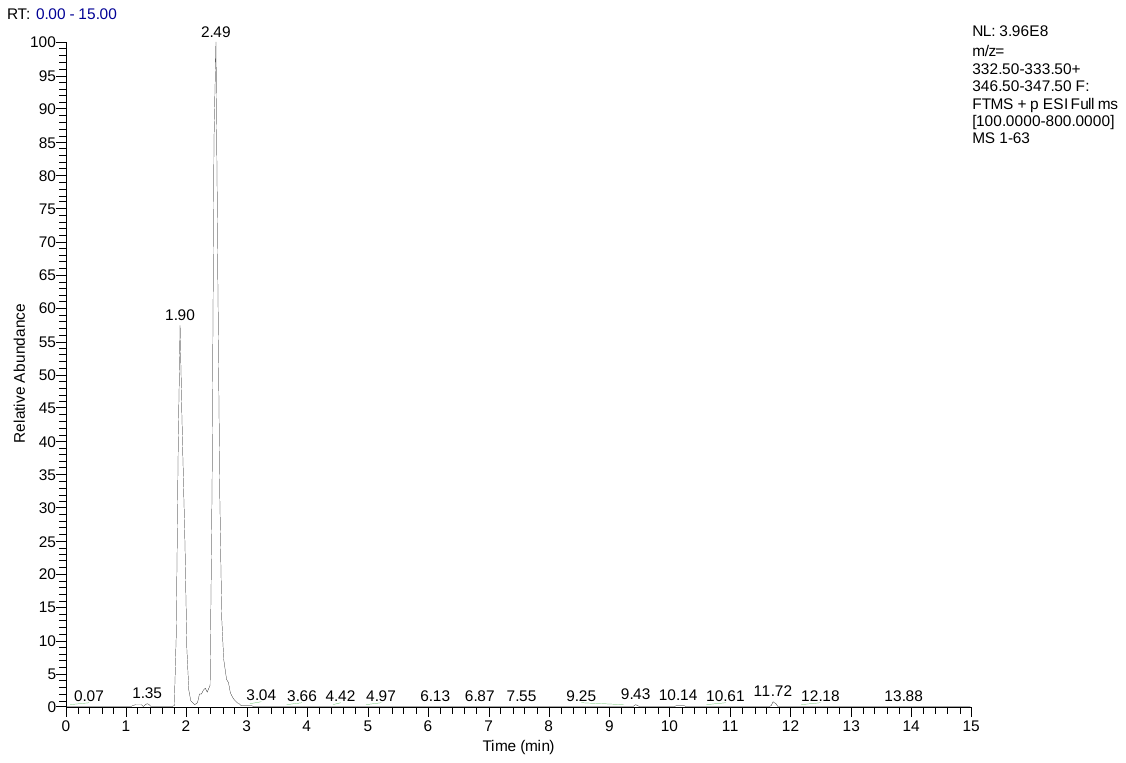

**S17. Oscillatoria nigro-viridis PCC 7112(L-S)**

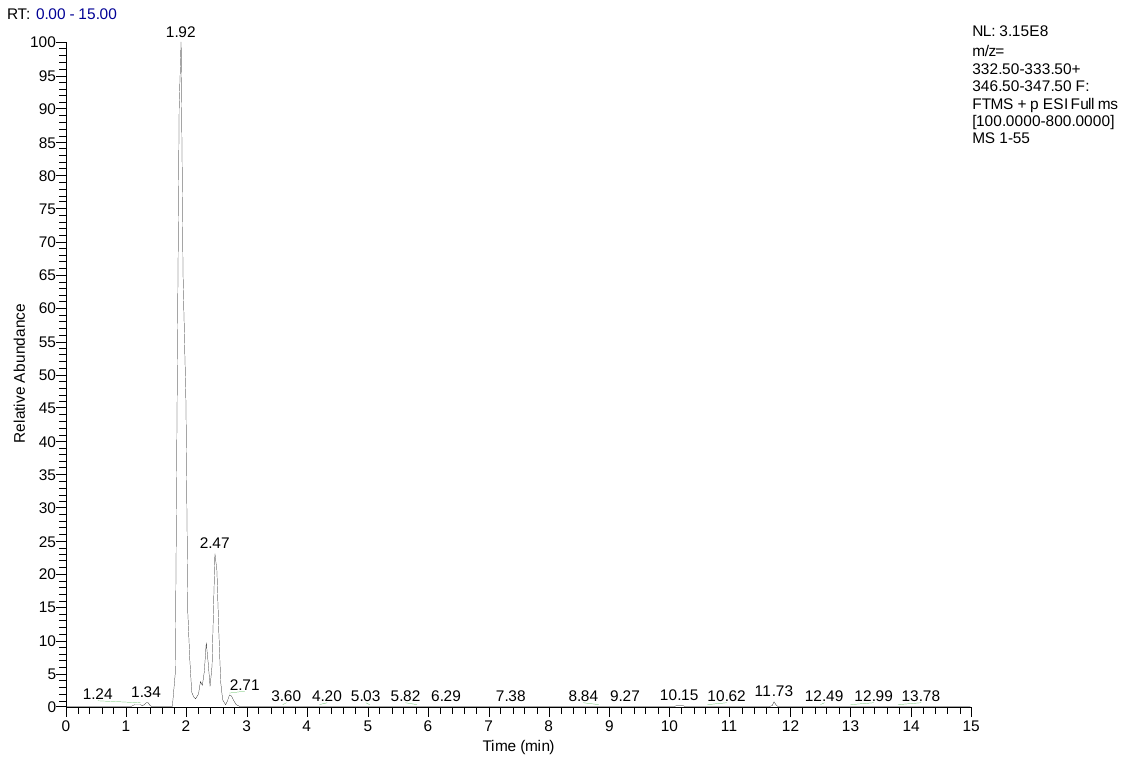

**S18. Gloeocapsa sp. PCC 73106(F-A)**

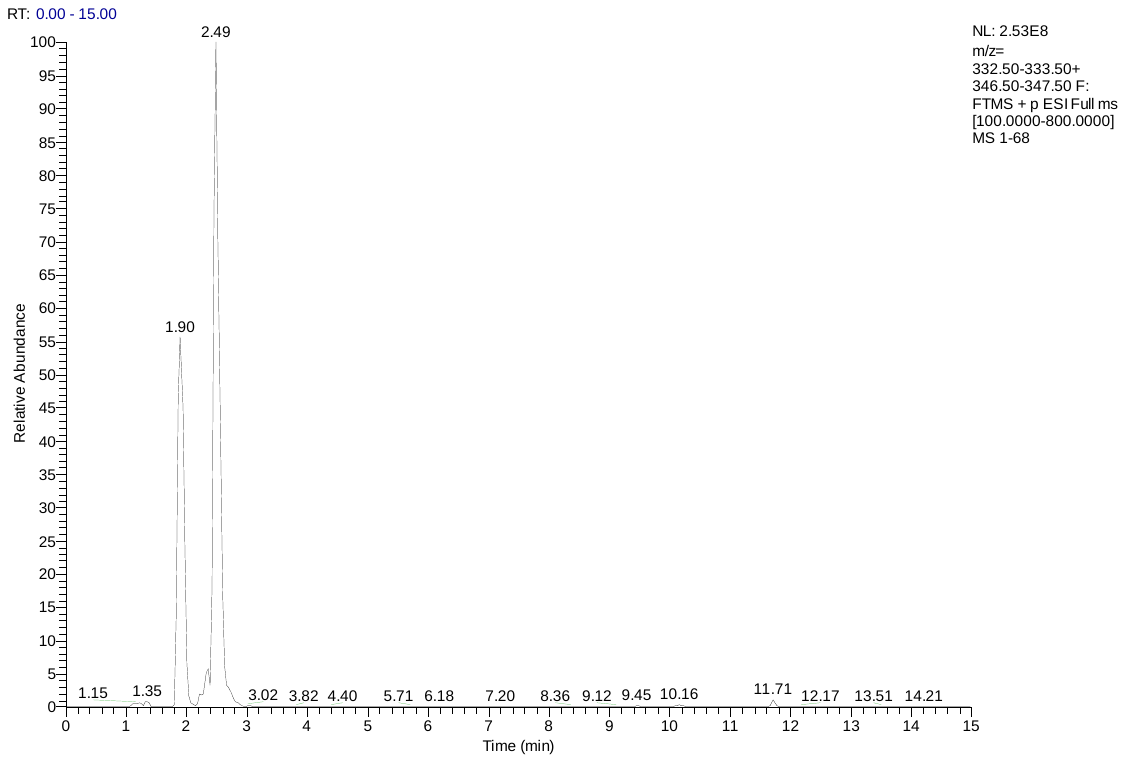

**S19. Microcystis aeruginosa PCC 7806(C-A)**

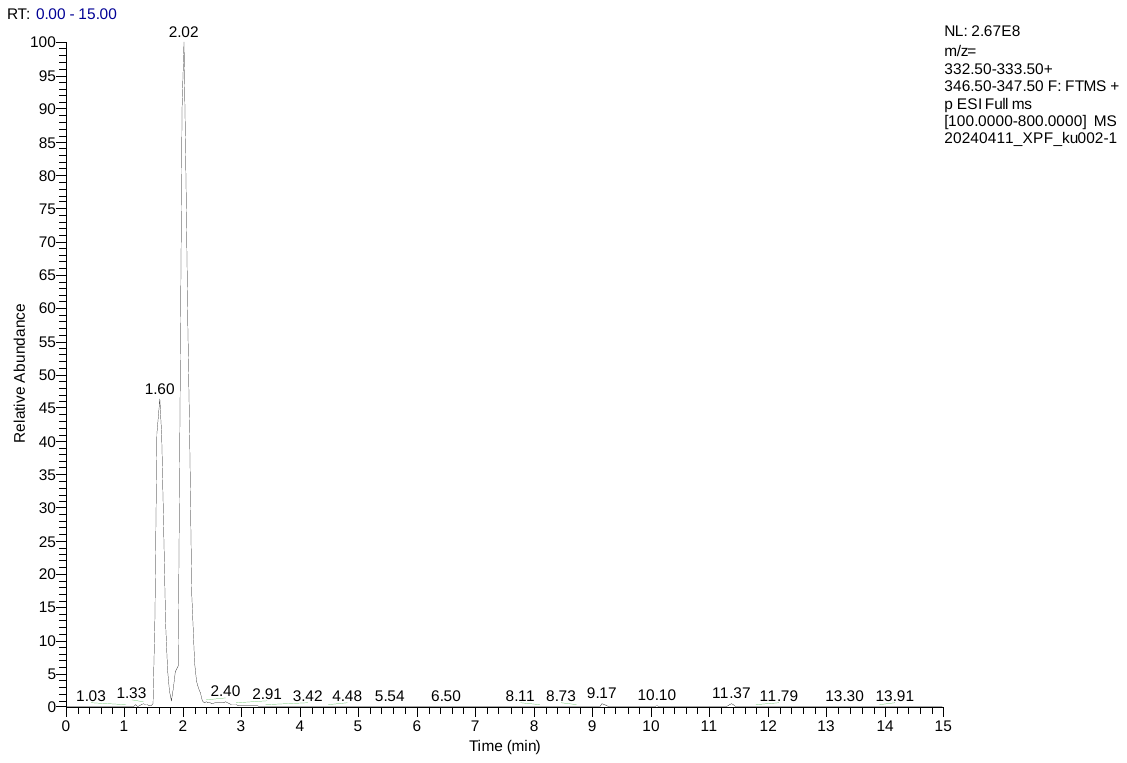

**S20. Coleofasciculus chthonoplastes PCC 7420(C-A)**

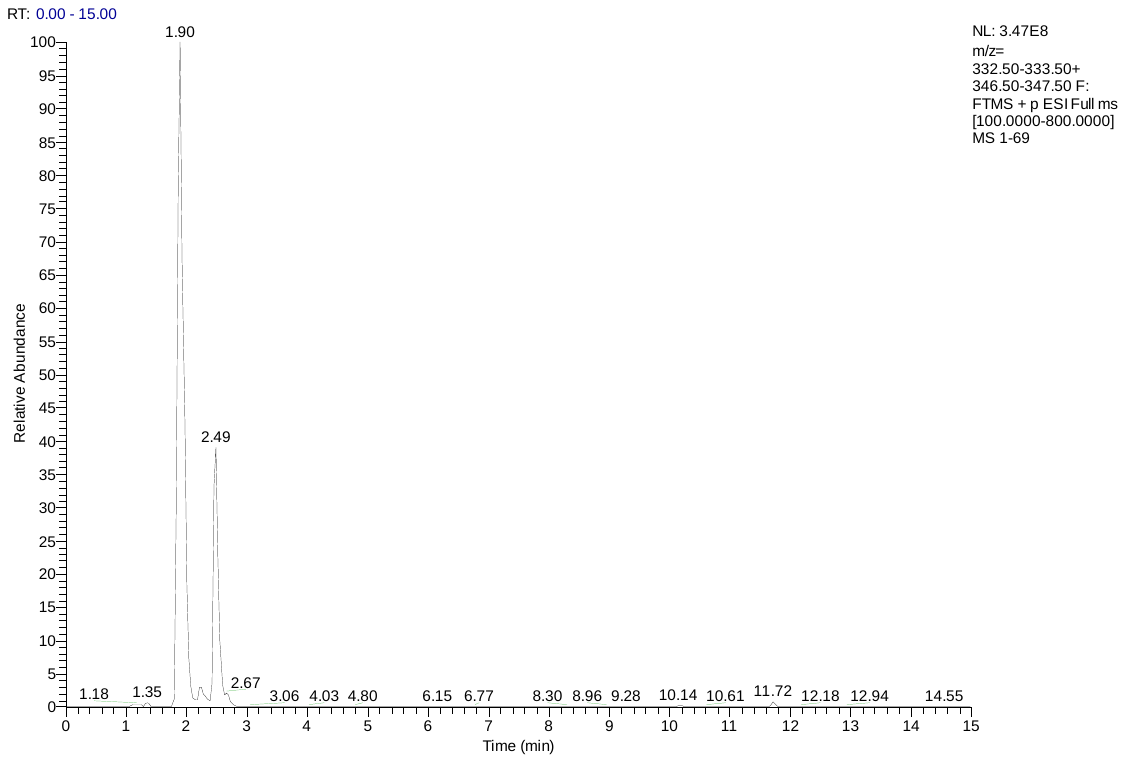

**S21. Cyanothece sp. BG0011(Y-A)**

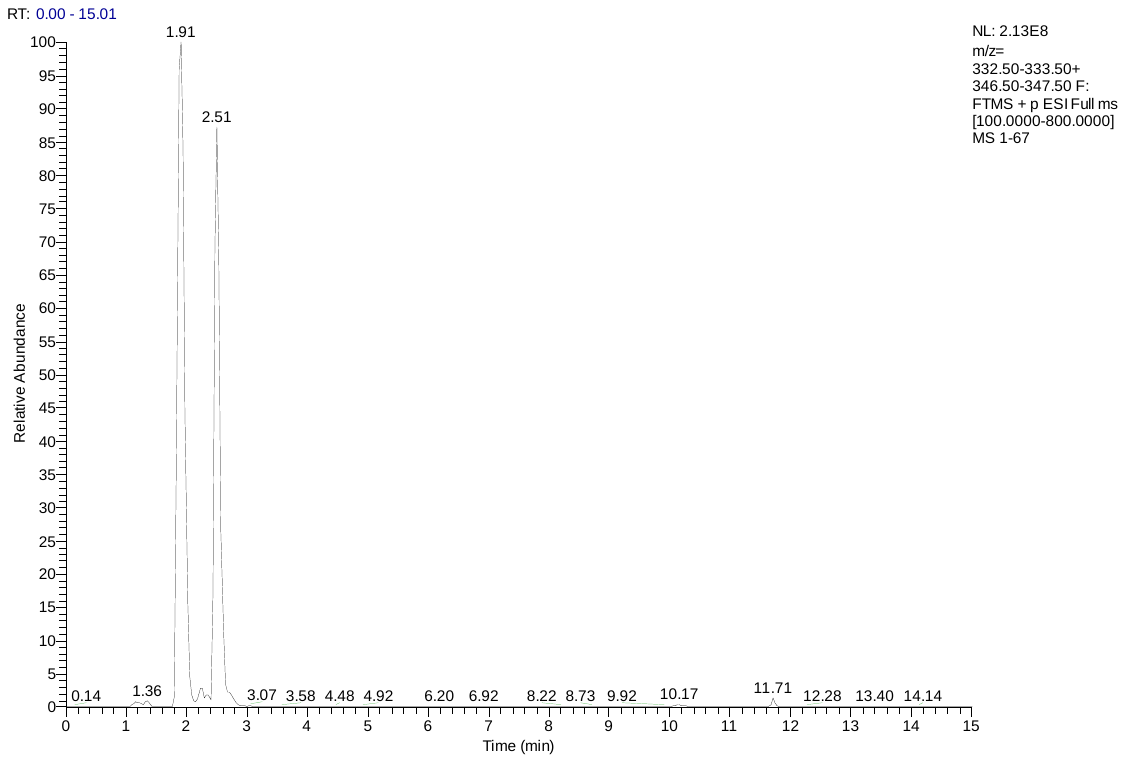

**S22. Crocosphaera chwakensis CCY0110(Y-A)**

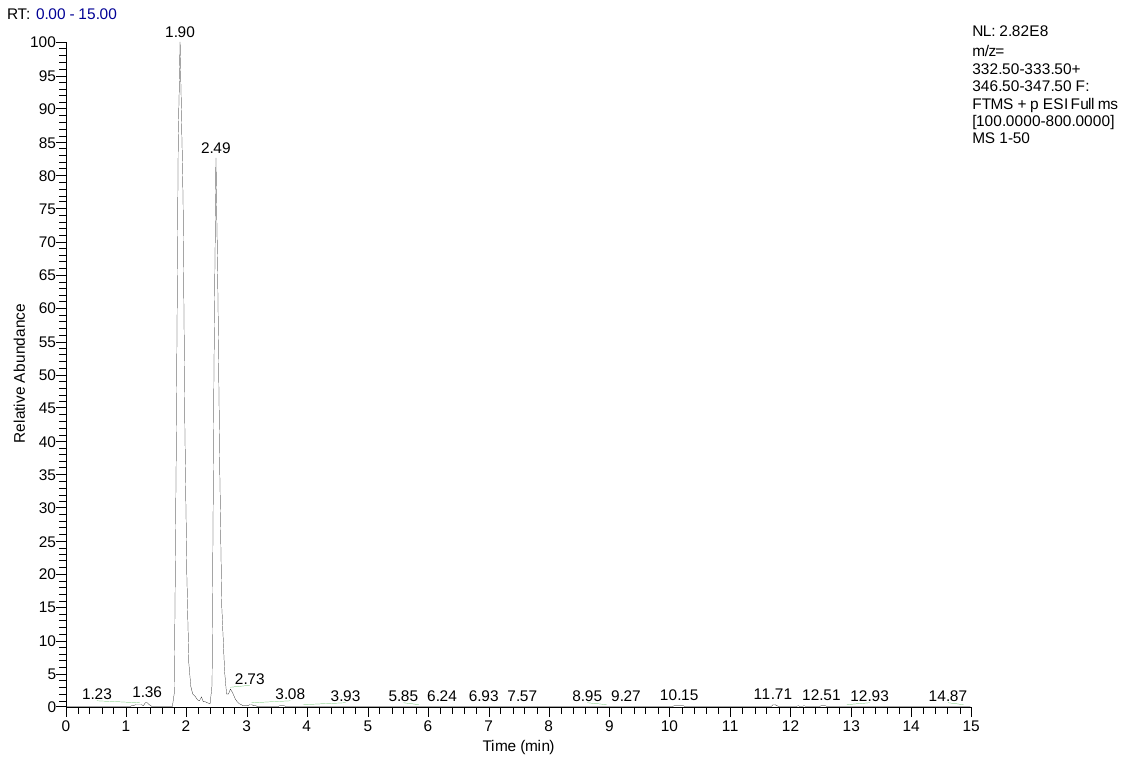

**S23. Nostoc sp. PA-18-2419(Y-S)**

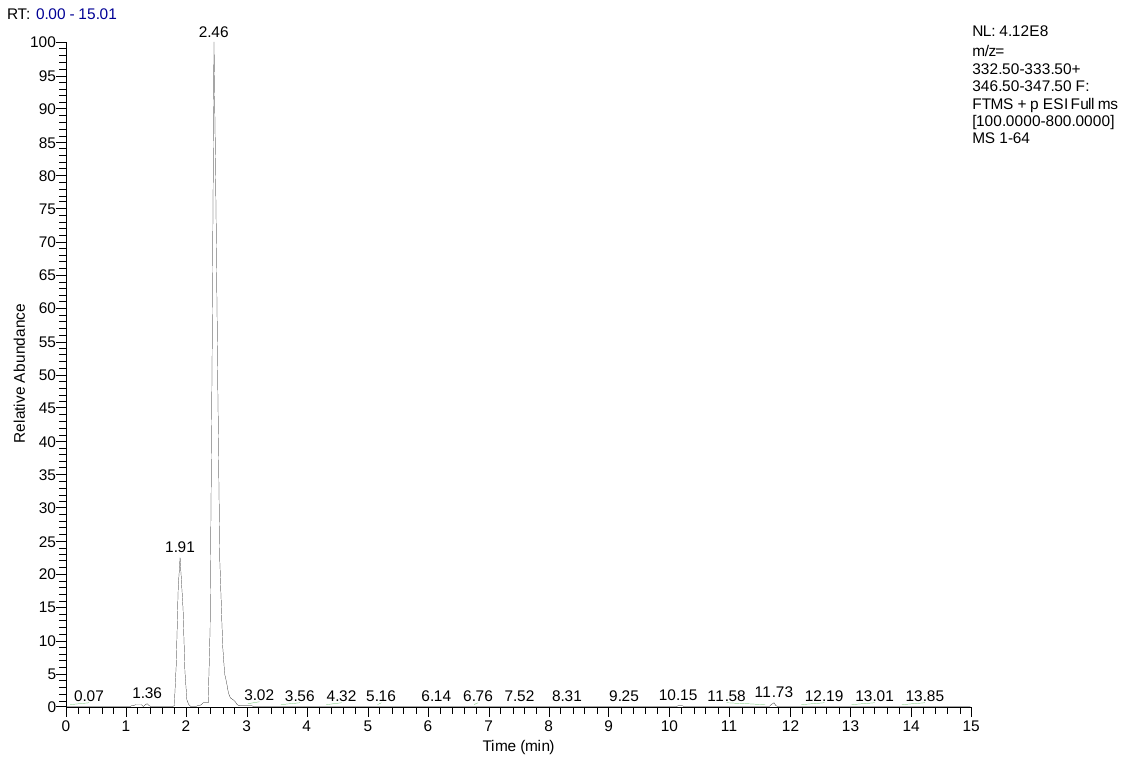

**S24. Nostoc sp. NIES-4103(Y-S)**

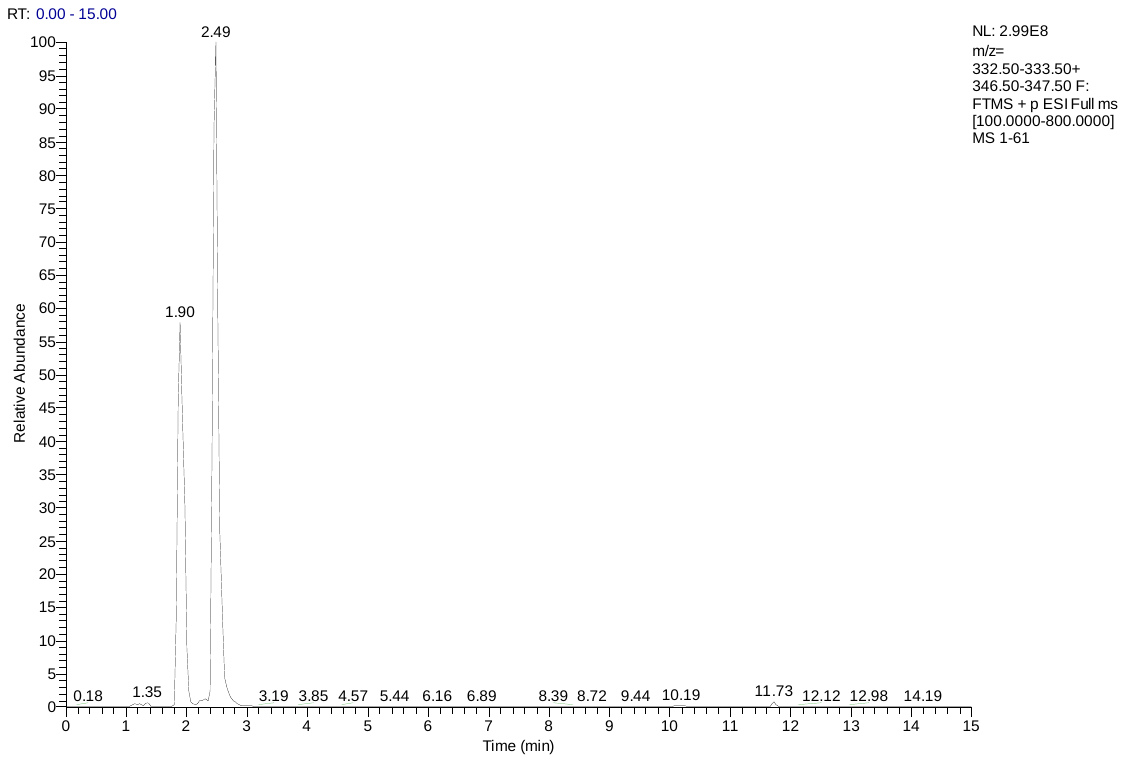

**S25. Calothrix elsteri CCALA 953(A-S)**

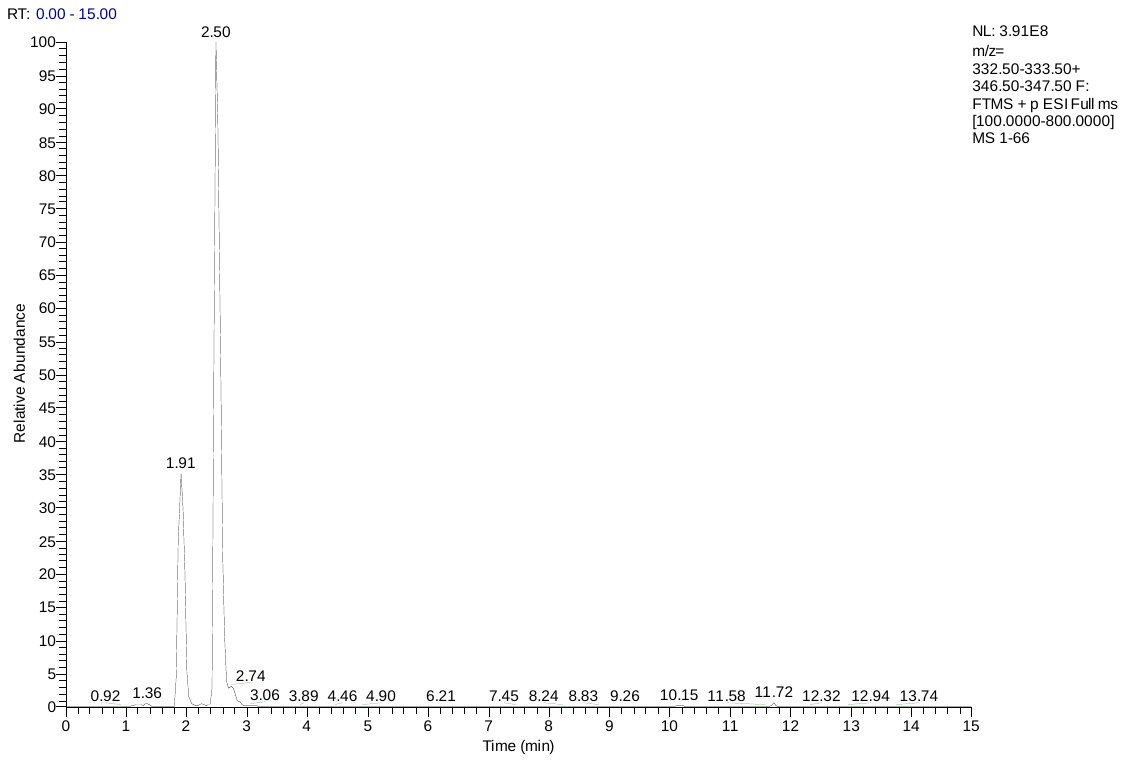

**S26. Calothrix sp. PCC 6303(A-S)**

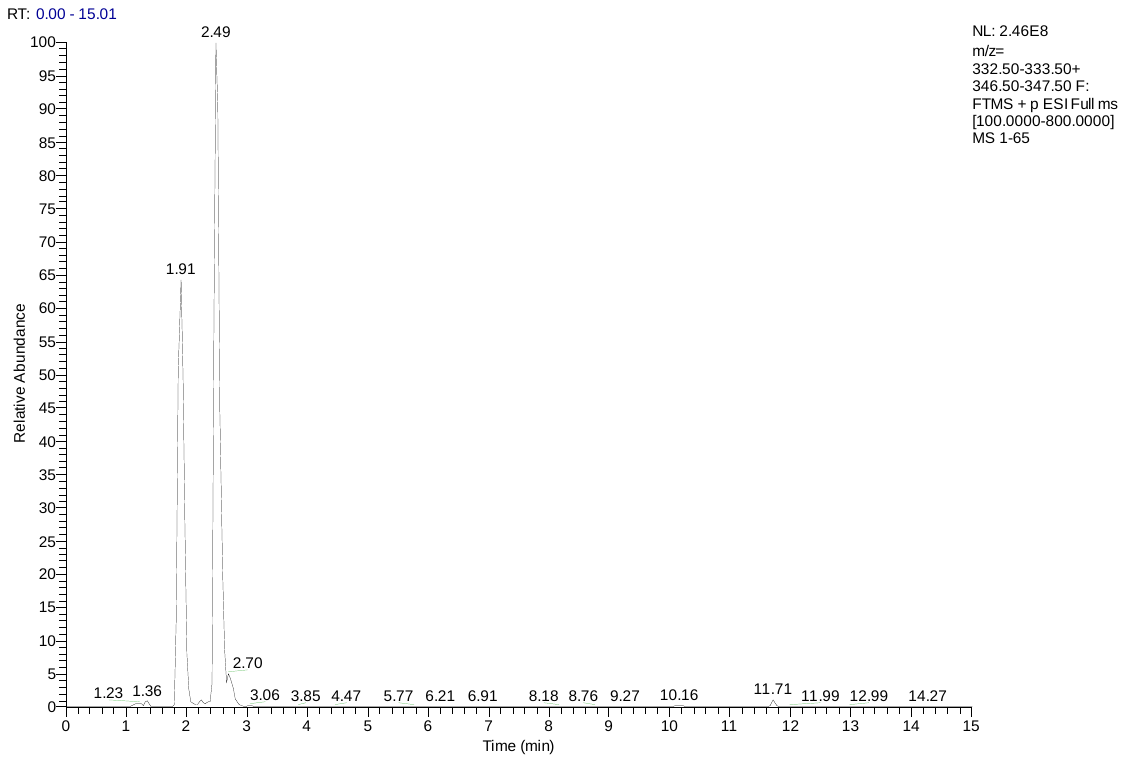

**S27. Chamaesiphon polymorphus CCALA 037(M-S)**

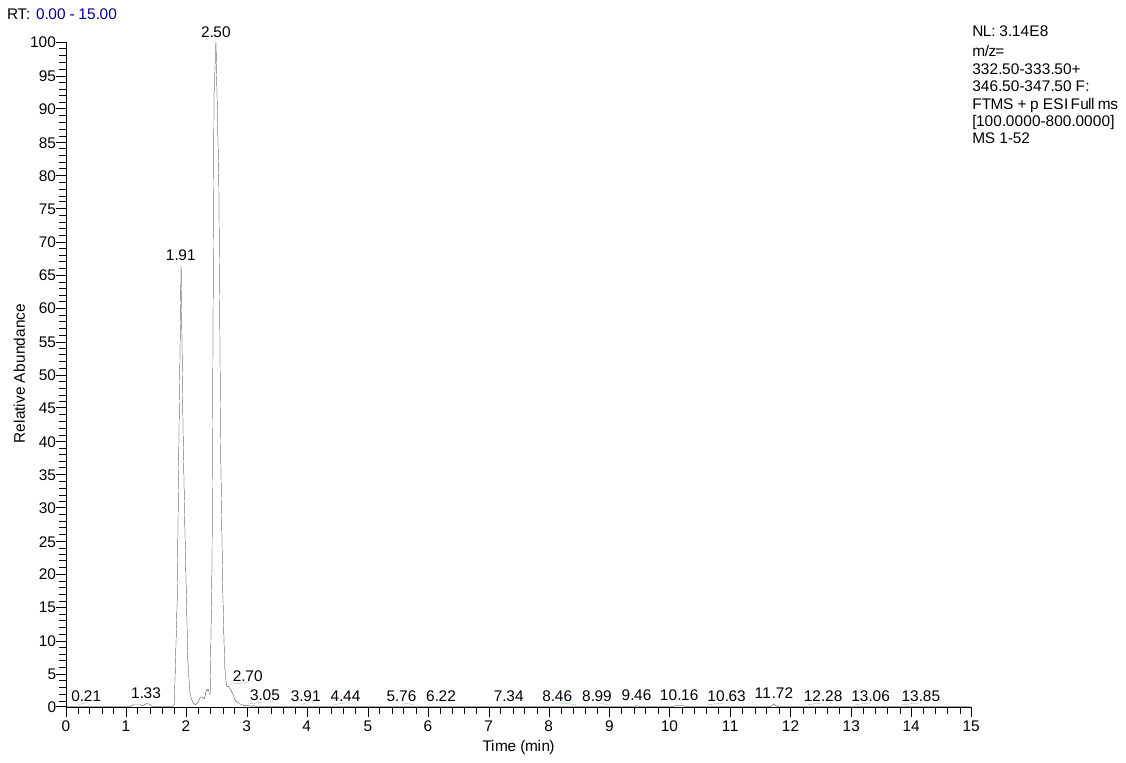

**S28. Chamaesiphon minutus PCC 6605(M-S)**

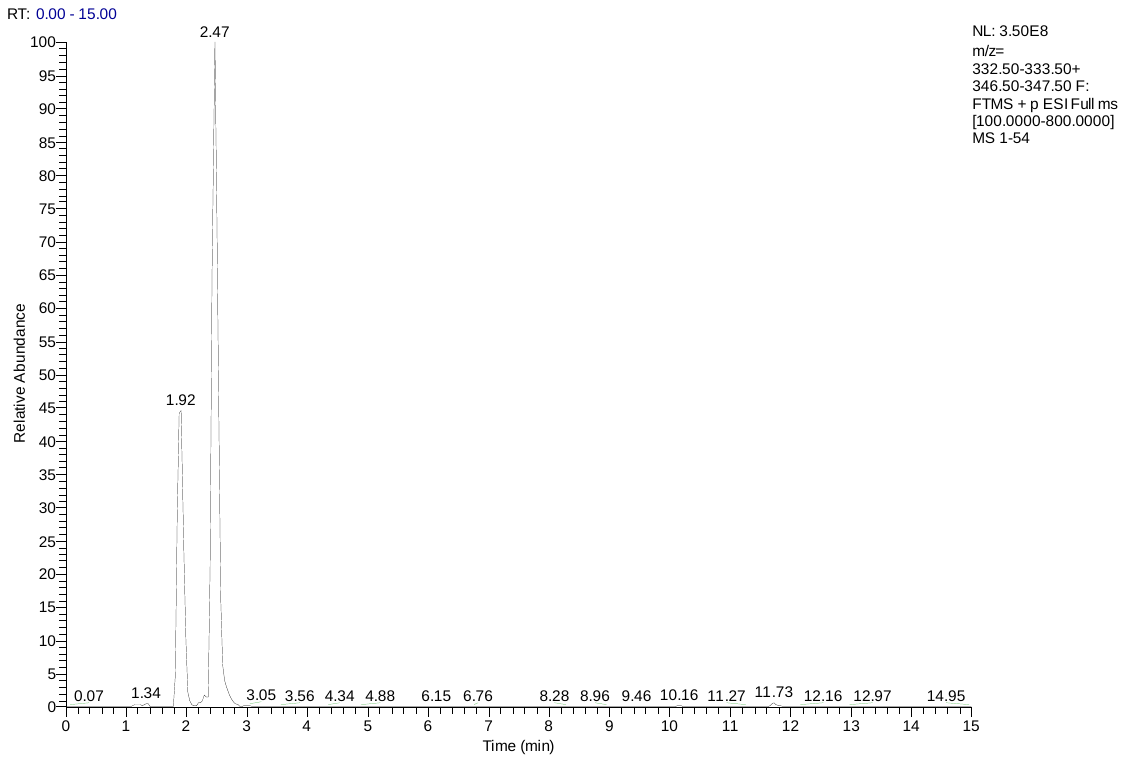
